# Per-sample standardization and asymmetric winsorization lead to accurate clustering of RNA-seq expression profiles

**DOI:** 10.1101/2020.06.04.134916

**Authors:** Davide Risso, Stefano M. Pagnotta

## Abstract

**Motivation:** Data transformations are an important step in the analysis of RNA-seq data. Nonetheless, the impact of transformations on the outcome of unsupervised clustering procedures is still unclear.

**Results:** Here, we present an Asymmetric Winsorization per Sample Transformation (AWST), which is robust to data perturbations and removes the need for selecting the most informative genes prior to sample clustering. Our procedure leads to robust and biologically meaningful clusters both in bulk and in single-cell applications.

**Availability:** The AWST method is available at https://github.com/drisso/awst. The code to reproduce the analyses is available at https://github.com/drisso/awst_analysis.

## 1 Introduction

Three major issues have caught most of the attention in the statistical analysis of gene expression data: (1) the classification or clustering of samples in molecularly-defined classes (e.g., subtypes of a disease; Dudoit and Fridlyand, 2002; Dudoit *et al*., 2002), (2) the identification of gene-level differences between groups of samples (i.e., differential expression; McCarthy *et al*., 2012), and (3) the functional characterization of the differentially expressed genes in pre-defined categories (i.e., gene-set enrichment analysis; Geistlinger *et al*., 2020).

Most of the statistical contributions in these areas focus on either the identification of data pre-processing transformations to remove systematic biases (i.e., normalization; Dillies *et al*., 2013), on the choice of the best inferential model, such as the negative binomial model (Robinson and Smyth, 2007), or on robust and flexible clustering techniques (Risso *et al*., 2018). Less attention has been paid to data transformations useful to improve the outcome of unsupervised clustering procedures. This is the focus of the present article.

Data pre-processing is often the first step before the application of downstream statistical methods. To the end of meeting as close as possible the assumptions of a statistical model, data transformations may be used. Prior to the unsupervised clustering of RNA-seq data, a logarithmic transformation and a standardization step are often performed. Recent workflows proposed in the literature (e.g., TCGA Research Network, 2015; Radovich *et al*., 2018) differ for the order in which the two steps are performed and for the estimation of the location and scale parameters for the standardization (see Supplementary Information). One of the expected effects of such procedures is to mitigate the influence of the outlying gene expressions, but, at the same time, the logarithm can increase the variance of the many lowly expressed genes. A selection of “informative genes” can ameliorate this issue empirically, but misuse of the log transformation can lead to erroneous results (Feng *et al*., 2014) and biases (Lun, 2018).

Most clustering procedures involve the computation of similarity/dissimilarity measures across samples and between clusters. Two recent papers (Jaskowiak *et al*., 2018; Vidman *et al*., 2019) evaluate the performance of clustering applied to complex RNA-seq datasets. They recommend (a) the average linkage for hierarchical clustering applied to correlation-based distances and (b) the PAM (Kaufman and Rousseeuw, 1990) procedure paired with the euclidean distance. On the other hand, applied studies (TCGA Research Network, 2015; Radovich *et al*., 2018) use the resampling methodology of Monti *et al*. (2003). In this context, too, average linkage and correlation distance, or PAM and euclidean distance, are common choices.

A critical point of any pipeline for clustering, and other procedures for omic-data, is feature selection. This general procedure often suffers from a subjective choice of the genes to include in downstream analysis. Specifically, in unsupervised clustering, the implicit assumption is that only the highest variable genes (typically 1,000–2,000) carry information about the samples mutual similarity. Furthermore, the choice of the variability index is also subjective.

Overall, there is a lack of clear guidelines on the optimal transformations and feature selection procedures for the clustering of RNA-seq data. In this landscape, our contribution is two-fold: (a) a data transformation that accounts for the count nature of the data and (b) a less subjective feature selection. In particular, we propose a two-step statistical transformation: (1) standardize the original data according to per sample estimates of location and scale; (2) smooth the standardized values with an asymmetric function that mimic winsorization in mitigating the effect of outliers. Our transformation applies independently to each sample; in this view, it is a per-sample transformation and we name it Asymmetric Winsorization per Sample Transformation (AWST). As for feature selection, we propose the use of Shannon’s entropy to remove uninformative genes.

## 2 Methods

AWST aims to regularize the original read counts to reduce the effect of noise on the clustering of samples. In fact, gene expression data are characterized by high levels of noise in both lowly expressed features, which suffer from background effects and low signal-to-noise ratio, and highly expressed features, which may be the result of amplification bias and other experimental artifacts. These effects are of utmost importance in highly degraded or low input material samples, such as tumor samples and single cells.

AWST comprises two main steps. In the first one, namely the *standardization step*, we standardize the counts by centering and scaling them, exploiting the log-normal probability distribution. We refer to the standardized counts as *z-counts*. The second step, namely the *smoothing step*, leverages a highly skewed transformation that decreases the noise while preserving the influence of genes to separate molecular subtypes. We define an asymmetric smoother, which is similar in spirit to winsorization (Tukey, 1962). The need for a differential treatment of expression values at the two tails of the distribution arises from the nature of the data: in fact, while the left tail is bounded by zero, the right tail depends on the amount of gene expression in the sample, the number of sequenced reads and the copy-number alterations, especially prevalent in cancer samples (Shao *et al*., 2019). We refer to the output of the two steps as the *smoothed-counts*. A further filtering method is suggested to remove those features that only contribute noise to the clustering.

### 2.1 Standardization step

Let us denote with *x_i,j_* the read counts for gene *j* in sample *i*. Several authors have noted that the right tail of log *x_i_* is akin to a Gaussian density (Hebenstreit *et al*., 2011; Hoyle *et al*., 2002; Gierliski *et al*., 2015; Okrah and Corrada Bravo, 2015) (Figure 1a). We therefore decide to anchor the standardization of the counts to this tail, by assuming that, for each sample *i, x_i_* is well approximated by a log-normal distribution (the dashed red curve in Figure 1a), i.e., *X_i_* = *e^μ_i_+σ_i_Z^*, where *Z* is a standard Gaussian random variable. Since our transformation is sample specific, we will omit the index *i* in the following, as it applies to any sample.

**Figure 1:**
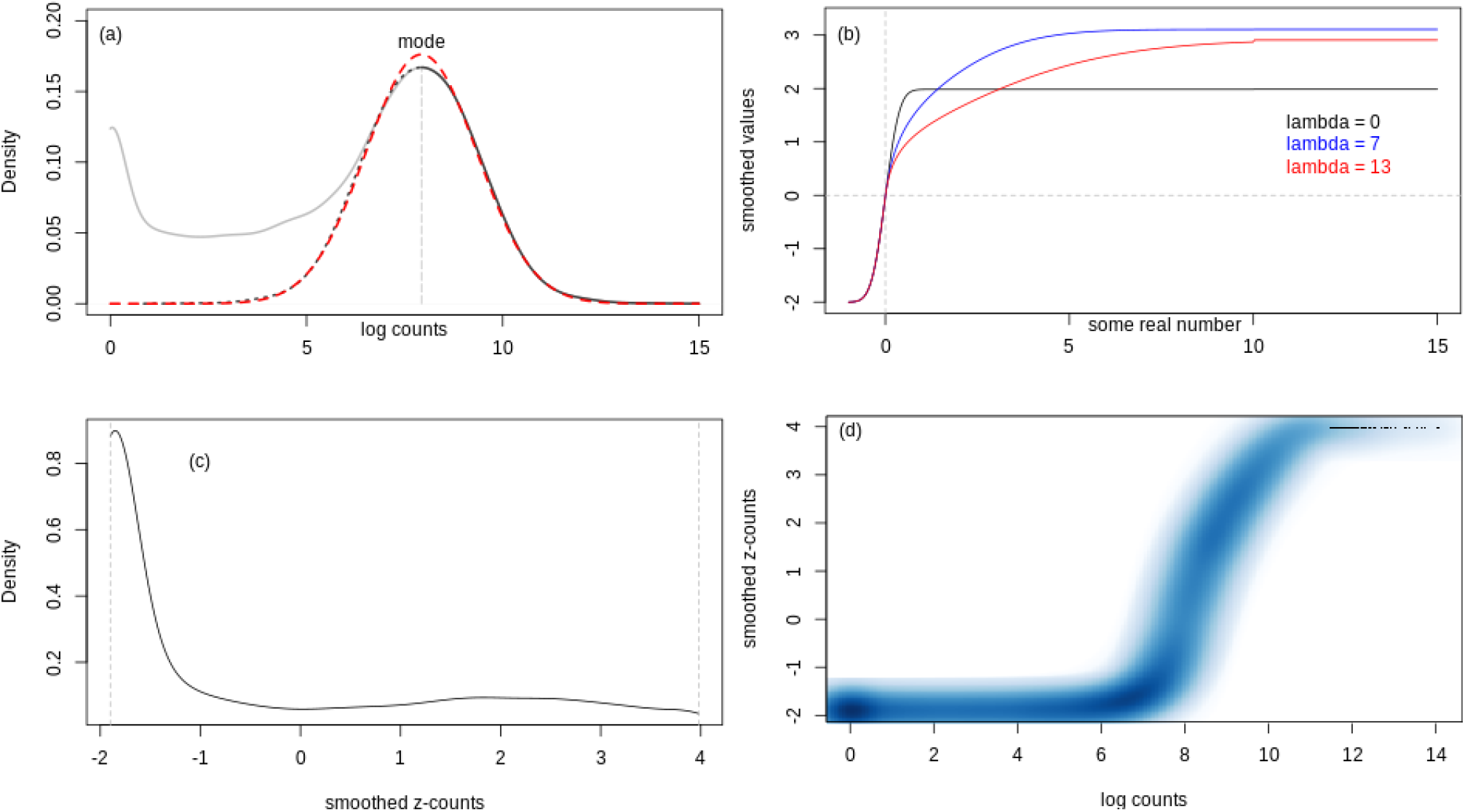
Illustration of AWST applied to sample TCGA-Q9-A6FU-01A-11R-A31P-31. (a) Kernel-density estimate of the probability density function (p.d.f.) of the log counts (grey curve). The dotted curve is the mirroring of the gray curve on the right of the mode. The red curve is the Gaussian p.d.f. with the location in the mode, and scale estimated from the values beyond the mode. (b) Graphs of the smoothing transformation with *σ*_0_ = 0.075 for λ = 0 (black curve corresponding to the Gaussian distribution function with location in 0 and *σ* = 0.075), λ = 7 (blue curve), and λ =13 (red curve). (c) Estimate of the p.d.f. of the smoothed z-counts (*σ*_0_ = 0.075, λ = 13). (d) Smoothed scatter-plot of the log counts versus their smoothed values (*σ*_0_ = 0.075, λ = 13).

For each sample, we first estimate *μ* with the right mode of the log2-counts empirical distribution, denoted as 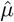. Analogously, we estimate the location 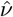 of the counts as 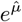. Then, we mirror the values on the right of the mode (the dotted blue curve in Figure 1a) and we estimate the variance 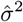 with the maximum likelihood estimator on the right tail and its mirrored values. To get the variance 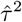 of the counts, we exploit the log-normal distribution, i.e., 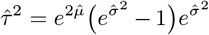. The standardized counts, the z-counts *z_j_*, are thus defined as

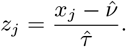

A similar approach was adopted by Hart *et al*. (2013) to standardize the log-FPKM values with the purpose of identifying a threshold that separates active from background genes. Both FPKM and the log transformation may lead to biases due to the influence of highly expressed genes (Bullard *et al*., 2010) and zero counts (Lun, 2018), respectively. Hence, we do not act on the log-FPKM transformed data. Rather, our standardization proposal is applied in the original (read count) scale. We shall note that our aim is different from that of Hart *et al*. (2013): while Hart *et al*. (2013) uses the transformation to classify between active and non-active genes, we use the transformation as a preliminary step for clustering.

### 2.2 Smoothing step

The second step of our proposal is a smoothing transformation applied to the *z-counts*. This transformation is crucial to regularize the expression of the genes affected by systematic artifacts that may bias the characterization of molecular subtypes. In fact, even when similar samples share the same subset of up-regulated genes, the level of up-regulation can change actively from sample to sample so that a compact cluster does not come to light. On the other hand, genes with a weak expression level contribute to the bias by acting as a source of noise for distance functions in high dimensions.

Briefly, we propose a smooth and skewed version of winsorization (Tukey, 1962) applied to the z-counts. To this end, we employ a smoothing function, *T*(·), based on a modified Gaussian distribution function, made asymmetric through the variance parameter. Let Φ(*z*) be the distribution function of a standard Gaussian random variable, and consider the function

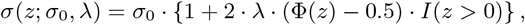

where *σ*_0_ is a constant and *I*(·) is the indicator function. *σ*(*z*; *σ*_0_, λ) = *σ*_0_ for all the *z* ≤ 0, and it is an increasing function for *z* > 0; the parameter λ ≥ 0 controls the growth-rate of the right tail. The asymmetrization mechanism follows the one adopted by Azzalini (1985) to define the skew-normal distribution. Differently from his proposal, we applied the asymmetrization device to the variance rather than to the body of the distribution.

The smoothing function is

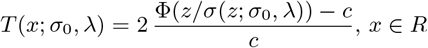

where

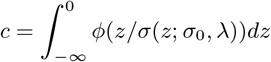

is a standardizing factor and *ϕ*(·) the probability density function of the standard Gaussian.

In Figure 1b, we plot the *T*(·) transformation for different values of λ. The black curve is the Gaussian probability distribution function with *μ* = 0 and *σ* = 0.075. The value of *σ* is empirically computed so that zeroes and counts close to zero are pushed to the minimum. As λ increases, from 7 (blue curve) to 13 (red curve), we observe a delay (as *x* increases) in transforming the values to the maximum. When λ = 0, the smoother is very close to acting as a device that dichotomizes the original values, where the mode is the changing point. Figure 1c, in which we draw the density estimates of the smoothed values of one TCGA sample, shows that the majority of the values have been shrunk around −2, while the others values gradually increase up to around 4. The effect of reducing the contribution of lowly expressed genes, and of the winsorization for the highly expressed ones, is made explicit by comparing the original log-counts versus the smoothed z-counts in Figure 1d. This latter graph shows that non-expressed and lowly expressed genes are uniformly transformed to the minimum, while the highly expressed genes are pushed to the maximum. The genes with an intensity around the mode (mapped to zero) can contribute with different levels to a distance function.

### 2.3 Gene filtering

The effect of the smoothing function is to push outlying genes to a maximum value and genes with a low expression level to a minimum. Genes reaching the maximum across all samples are irrelevant, as well as those always assigned to the minimum value. Moreover, the features with about the same expression level can only add noise to the distance between samples without contributing meaningfully to the similarity measure. To remove the irrelevant features, and those only contributing noise, we propose a filtering procedure based on a heterogeneity index, which is possible only because AWST forces the expression level of any sample in the same range.

Since the expression levels for every sample are bound by the same maximum and minimum values, we define a partition of *K* equally spaced intervals. The partition can be used to categorize the gene expression levels in each sample, and then to compute the heterogeneity index *h_j_* for each of the genes. Given a cutoff value *c*, we remove all those features fulfilling the constraint *h_j_* < *c*. As a heterogeneity index we use the normalized Shannon’s entropy

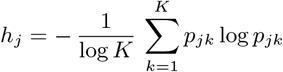

where *p_jk_* is the empirical probability associated with the *k^th^* interval of the partition and the *j^th^* gene.

Our filtering procedure maps the samples in a less parsimonious subspace than that induced by the rule of thumb of retaining the top 1,000 features (or few more), but it often yields a finer partition of the samples thanks to the AWST, as we show in section 3.

### 2.4 Parameters choice

Our transformation has two tuning parameters: λ and *σ*_0_, which control the amount of smoothing. Larger λ values push fewer points to the maximum value (Figure 1). Similarly, larger *σ*_0_ values lead to less shrinkage. Empirically, we found that values of λ ranging from 5 to 13 are good choices in practice. The default value of *σ*_0_ is chosen to shrink to the minimal value the low counts, more likely to be noise, while simultaneously preserving a dynamic range of moderate and high values.

The two parameters both contribute to the amount of shrinkage. Decreasing λ increases the shrinkage for highly expressed genes, while decreasing *σ*_0_ increases the overall rate of shrinkage, eventually leading, for very small values of *σ*_0_, to dichotomization of the data. In all the analyses of this article, we set the two parameters to their default values (*σ*_0_ = 0.075, λ = 13).

Similarly, our filtering step has two tuning parameters: the threshold *c* and the number of bins *K*. The threshold *c* controls the amount of genes that get filtered out according to the normalized entropy. An entropy of 0 indicates the genes that will not contribute any information in the classification, falling in the same bin across all samples. In practice, a small positive value (we adopt 0.1 by default) may be preferable to 0, to additionally remove those genes that contribute very little to the classification procedure.

Also the number of bins *K* contributes to the number of features retained for downstream analyses. We empirically found 20 as a good value, being a compromise between the retention of informative features and the removal of the non-informative ones. As *K* decreases, the intervals become larger and a higher number of genes is removed; on the contrary, when *K* increases the filtering procedure retains more genes with a possible increase of noise in downstream analyses.

## 3 Results

### 3.1 Synthetic data

First, we evaluate our procedure on a synthetic dataset created from the SEQC (Su *et al*., 2014) reference data. Briefly, starting from 80 real RNA-seq technical replicates of Agilent Universal Human Reference, we simulate differential expression to create five groups of samples and we compare each procedure based on its ability to retrieve the simulated true partition (see Supplementary Information for details on how differential expression was simulated). Our synthetic dataset consists of 150 samples and 19,701 genes. The AWST procedure yields the same results both with and without our gene-filtering step. We present the results omitting the gene filtering, i.e., without removing any of the 19,701 genes.

The first set of experiments compares AWST versus a pre-processing method inspired by the transformation of Hart *et al*. (2013) and state-of-the-art procedures proposed for the clustering of RNA-seq data in Radovich *et al*. (2018) and TCGA Research Network (2015). Radovich and TCGA pre-processings (detailed in Supplementary information) adopt almost the same robust pipeline; they differ for the choices of the robust estimates of location and scale, and for the order in which standardization and log2-transformation are performed (see Supplementary Information). We also include in the comparison a simpler procedure that retains the top variable genes after FPKM normalization. The comparisons are two-fold: a) we apply hierarchical clustering (with Euclidean distance and Ward’s linkage) to each pre-processing and evaluate the agreement between the inferred partition and the true clusters using the Adjustud Rand Index (ARI) (Hubert and Arabie, 1985); b) with the help of silhouette plots (Rousseeuw, 1987), we measure the compactness of the true partition in the distance matrices obtained after each pre-processing.

Figure 2 shows the dendrograms of hierarchical clustering with Euclidean distance and Ward’s linkage applied to each pre-processing. Each tree is paired with the Calinski and Harabasz (1974) (CH-) curve from the cophenetic matrix. The CH-curve is essentially the sequence of the ANOVA test-statistics varying the number of groups from the dendrogram. Local maxima, or even better a global maximum, suggest the number of clusters in the data.

**Figure 2.**
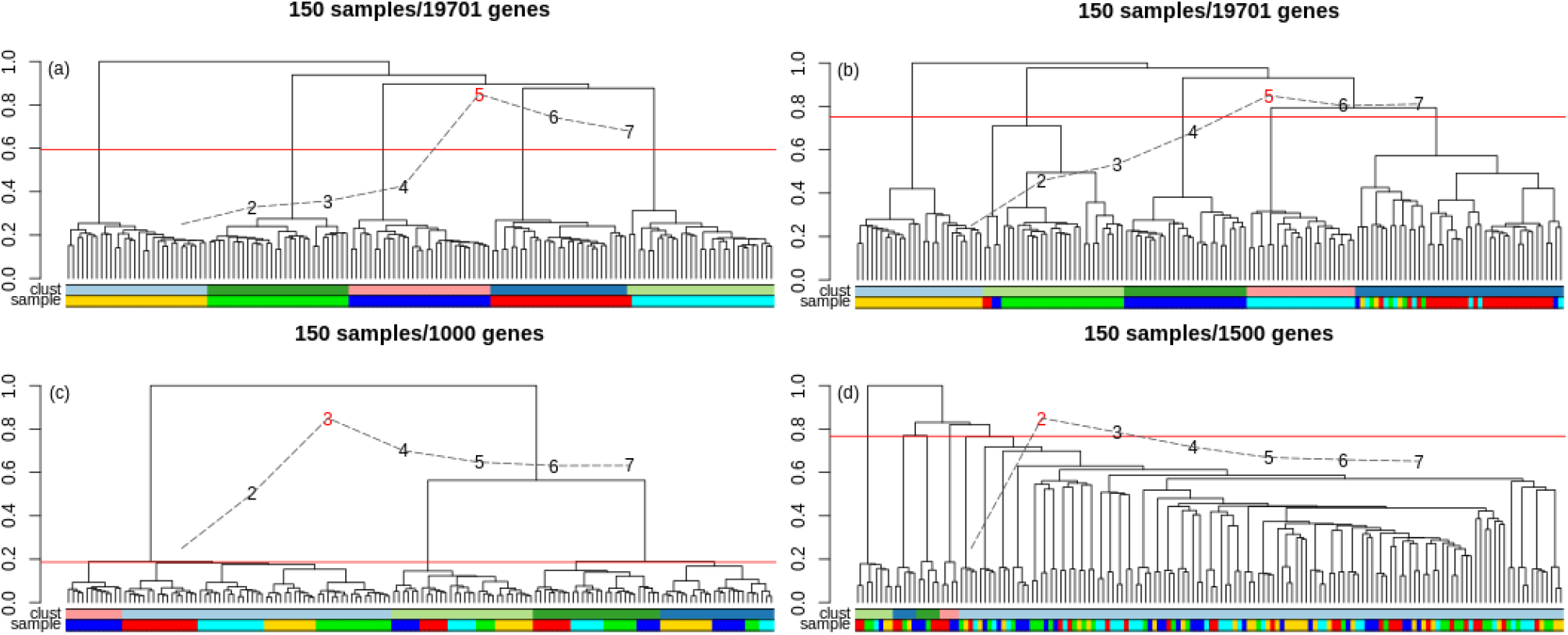
Performance of different data pre-processing paired with the hierarchical clustering (Euclidean distance, Ward’s linkage). a) AWST data pre-processing; b) Hart’s data pre-processing; c) Radovich’s data preprocessing; d) TCGA data pre-processing. The “sample” bar indicates the true partition, and the “clust” bar indicates the inferred partition obtained by cutting the tree to get 5 clusters. The Calinski-Harabasz curve is superimposed in each panel.

Our AWST procedure leads to a quasi optimal clustering (ARI = 0.98; Figure 2a); a slightly worse result is obtained with Hart pre-processing (ARI = 0.68; Figure 2b). The pre-processings of Radovich (ARI = 0.08; Figure 2c) and TCGA (ARI = 0.001; Figure 2d) clearly do not recover the true partition. A simpler procedure that selects the top 2,500 features after FPKM normalization works better (ARI = 0.46; Supplementary Figure 1e). However, such procedure is inherently sensitive to the number of selected genes. In fact, it fails when we double the amount of features (ARI = 0.001; Supplementary Figure 1f). We observe that the performance of the FPKM pre-processing (measured by ARI) decreases as we increase the number of features (Supplementary Figure 5, black line). Furthermore, AWST yields the highest average silhouette width (0.25; Supplementary Fig. 2a), confirming that our proposal preserves relevant information about the true partition in the data.

In the second set of experiments, we compare the four pre-processings with a state-of-the-art clustering strategy for complex RNA-seq experiments, i.e., the resampling methodology of Monti *et al*. (2003), implemented in the *ConsensClusterPlus* (Wilkerson and Hayes, 2010) R package. Both Radovich *et al*. (2018) and TCGA Research Network (2015) obtain their final consensus clustering by using *ConsensClusterPlus*.

Briefly, the consensus clustering (referred to as *outer clustering*) is a hierarchical clustering procedure with average linkage based on 1,000 *inner hierarchical clusterings*, also with average linkage. Each time 80% of the original samples are randomly drawn. Both papers adopt Pearson’s correlation between samples as a distance matrix. We applied the same method to each pre-processing.

When AWST is applied prior to the consensus clustering procedure, the approach leads to a perfect partition (Figure 3a). Hart transformation leads to a slightly less accurate partition (Figure 3b, ARI = 0.70) and, more importantly, to more uncertainty in the cluster allocations (as shown by the consensus matrix), suggesting that AWST improves the robustness of the clustering results. The Radovich procedure also yields a perfect partition (Figure 3c). However, that is not the case when, after Radovich pre-processing, we apply a simple hierarchical clustering (Figure 2c) or a different consensus clustering procedure based on Euclidean distance and PAM (Supplementary Figure 4c, ARI = 0.12), suggesting that AWST is more robust to the choice of the downstream clustering method. Conversely, the TCGA procedure results in a poor partition with large cluster uncertainty, both when the consensus clustering is based on the average linkage with Pearson’s correlation (Figure 3d, ARI = 0.09) and when it is based on the Euclidean distance and PAM clustering (Supplementary Figure 4d, ARI = 0.001).

**Figure 3.**
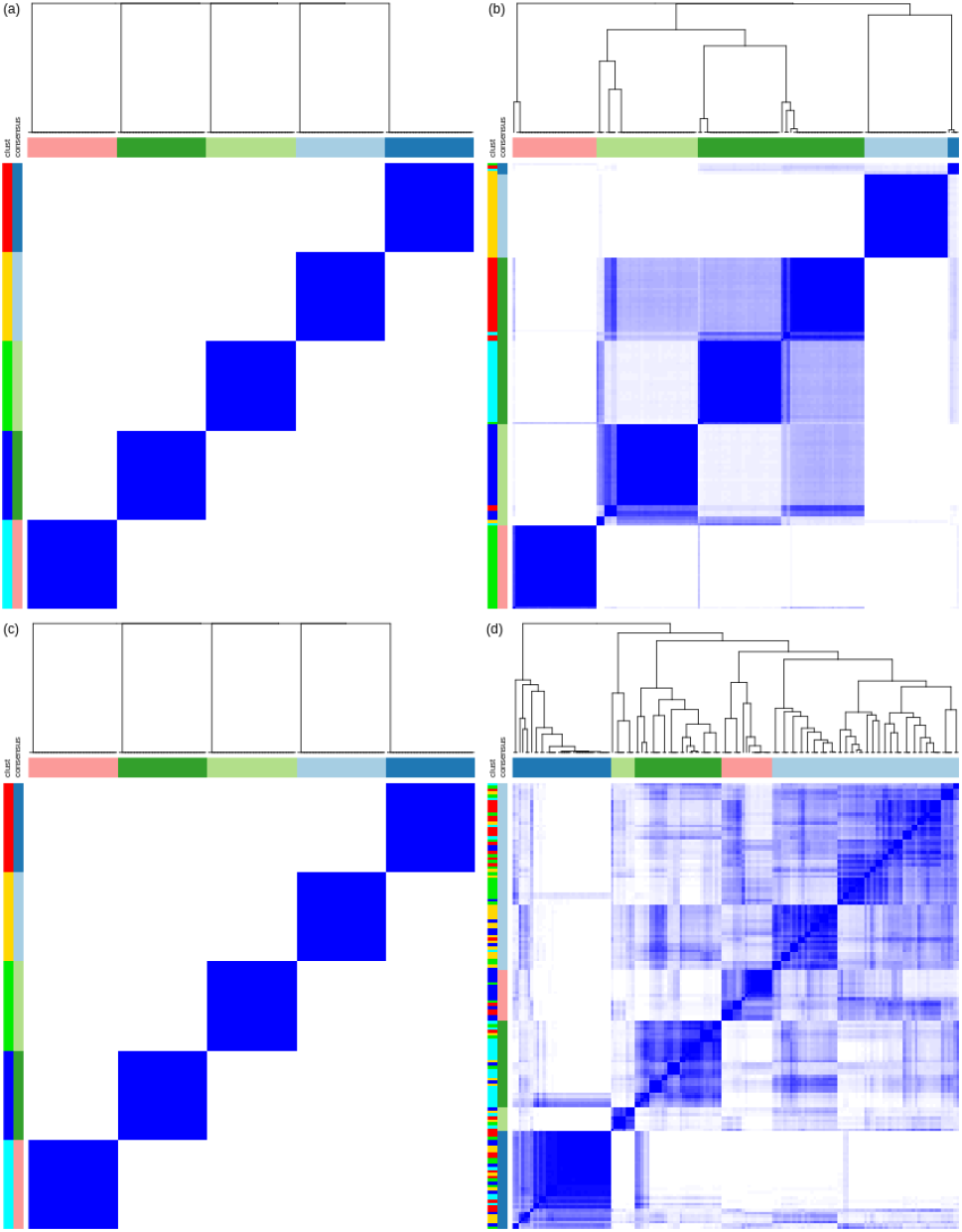
Performance of different data pre-processing paired with ConsensusClusterPlus (inner and outer average linkage with Pearson’s correlation as distance matrix). a) AWST data pre-processing; b) Hart’s data pre-processing; c) Radovich’s protocol; d) TCGA data protocol. The “clust” bar indicates the true partition, and the “consensus” bar indicates the inferred partition obtained by requiring 5 clusters.

The consensus clustering analysis confirms our observations on the FPKM pre-processing: the method works with the top 2,500 genes (Supplementary Figure 3e, ARI = 0.52), but completely fails when increasing this number to 5,000 (Supplementary Figure 3f, ARI = 0.18). The FPKM pre-processing with 2,500 genes works better when paired with the Euclidean distance and the PAM method (Supplementary Figure 4e, ARI = 1); the performance decreases when we consider 5,000 features (Supplementary Figure 4f, ARI = 0.61). Independently of the clustering procedure, the ARI decreases with the increase in the number of features (Supplementary Figure 5), indicating that selecting the correct number of most variable genes (an unknown quantity in real data) is paramount to the correct clustering of the samples.

Taken together, these results show that AWST is the only pre-processing that leads to an optimal sample partition on all the tested clustering algorithms (Figure 2a, Figure 3a, Supplementary Figure 4a).

### 3.2 Lower Grade Glioma (LGG) study

A collection of 516 primary tumor samples of the Lower Grade Glioma (LGG) study from The Cancer Genome Atlas (TCGA) allows to demonstrate the advantages of AWST on a complex RNA-seq experiment. We compare AWST results with those previously published, in which ad-hoc and sometimes multi-omic approaches were used. While there is no ground truth in this dataset, comparing our partition to those identified by several independent, multi-omic approaches, allows us to check if the derived clusters represent biologically meaningful subtypes.

LGG is classified in three molecular subtypes connected to DNA lesions (“Original subtype” in the plots). The first difference concerns the Isocitrate DeHydrogenase genes 1 and 2 (IDH) status. The wildtype of both genes defines an autonomous molecular subtype (IDHwt); in case of mutations, the co-deletion of chromosomes 1p/19q discriminates between CODEL (IDHmut-codel) and NON-CODEL (IDHmut-non-codel) molecular subtypes (Eckel-Passow *et al*., 2015). Ceccarelli *et al*. (2016) showed that the IDHmut-non-codel molecular subtype is equivalent to the G-Cimp subtype of Noushmehr *et al*. (2010).

We apply AWST to the RSEM estimated counts after GC-content correction and full quantile normalization. The gene-filtering step retains 10,212 genes out of the original 19,138. We compute the Euclidean distance followed by hierarchical clustering with Ward’s linkage (Figure 4). The lack of meaningful clusters in the dendrogram of the 8,926 filtered out genes (Supplementary Figure 11) indicates that the removed features only contribute noise.

**Figure 4.**
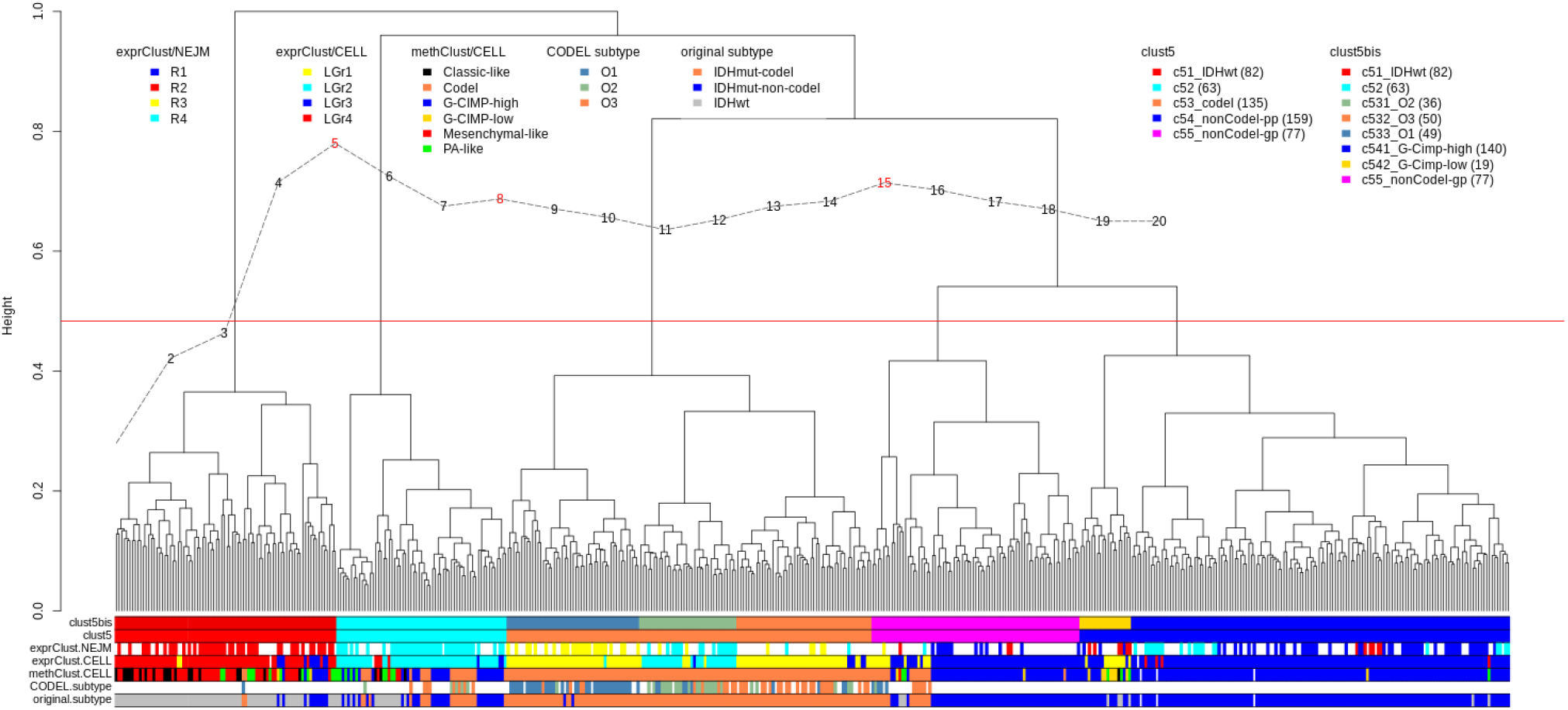
Hierarchical clustering of the 516 Lower Grade Glioma (LGG) samples with Ward’s linkage and Euclidean distance, involving 10,363 genes having normalized entropy greater than 0.10. The dashed curve is the Calinski and Harabasz index. The red horizontal line represents the selected resolution of the clustering, as suggested by the index. The color bands report, in addition to our cluster labels, state-of-the-art classification of the samples. clust5: our partition in 5 groups; clust5bis: our partition in 8 groups; exprClust/NEJM: partition obtained from expression data in TCGA Research Network (2015); exprClust/CELL: partition obtained from expression data in Ceccarelli *et al*. (2016); methClust/CELL: partition from methylation data in Ceccarelli *et al*. (2016); CODEL subtype: sub-classification of CODEL subtype defined in Kamoun *et al*. (2016); original subtype: Isocitrate DeHydrogenase genes 1 and 2 (IDH) wild type (IDHwt), co-deletion event of the chromosomes 1p/19q (IDHmut-codel); non-co-deletion (IDHmut-non-codel).

The dendrogram in Figure 4 is annotated with different partitions. The expression clusters denoted by the “exprClust/CELL” bar come from a pan-glioma study, in which stringent filtering was applied to select 2,275 features in a set of 667 samples (154 GBM and 513 LGG) (Ceccarelli *et al*., 2016), leading to four distinct groups. A second expression clustering (“exprClust/NEJM”) is from TCGA Research Network (2015) and provides a different partition in four groups. Kamoun *et al*. (2016) studied 156 oligodendroglial tumors of the IDHmut-CODEL subtype. With a multi-omic approach, they found three molecular subtypes (indicated by the “CODEL/subtype” bar in Figure 4).

The CH-curve suggests a primary partition (“clust5”) of 5 groups that mostly mirrors the Original subtype of brain cancer: c51 is the cluster of the IDHwt samples; c53 includes the IDHmut-codel samples. The IDHmut-non-codels splits into two groups: c54 and c55. Cluster c52 mostly corresponds to the LGr2 (Ceccarelli *et al*., 2016) and R4 groups (TCGA Research Network, 2015); the genomic determinants of this group are still unexplored. Clust5 is supported by the study of the overall survival time associated with each sample (Supplementary Figure 6; log-rank test *p*-value of ≤ 2.2 · 10^-16^).

C53 is a large collection of IDHmu-codel samples. Kamoun *et al*. (2016) specifically studied this kind of tumors and obtained three sub-groups, supported by different genomic analyses. Our hierarchical clustering also splits c53 into three sub-groups that match the corresponding ones of Kamoun. To support the identification of our sub-groups, we estimated the survival curves associated with them (Supplementary Figure 8; log-rank test p-value of 0.02). Our finding is comparable to Supplementary Figure 8a of Kamoun *et al*. (2016).

C54 is an almost homogeneous cluster of G-Cimp-high samples. Our clustering identifies c541 and c542. The latter matches the G-Cimp-low sub-type discovered in Ceccarelli *et al*. (2016), obtained using a supervised analysis of methylation data. Our experiment identifies the G-Cimp-low solely based on the expression profiles, for the first time to the best of our knowledge (Supplementary Figure 9; log-rank test *p*-value = 2 · 10^-4^).

To compare the performance of AWST to its competitors, we apply the other methods to the LGG dataset. None of the estimated partitions of the 516 LGG samples consistently match the subtypes known from the literature. (Supplementary Figures 12, 14, 16). Reassuringly, we note that our implementation of the TCGA protocol reproduces the original results of TCGA Research Network (2015), denoted in the figures by “exprClust/NEJM”.

### 3.3 Cord Blood Mononuclear Cells

To further demonstrate the performance of AWST, we apply the transformation to the Cord Blood Mononuclear Cells (CBMC) single-cell experiment of Stoeckius *et al*. (2017) used to introduce the CITE-seq methodology. Briefly, CITE-seq allows independently profiling of scRNA-seq and a panel of antibodies for each cell. The antibody-derived tags data (ADTs) measure the levels of those membrane proteins able to classify each cell to the sub-populations of the immune system, essentially providing the same informative content of flow cytometry. As an example, the combination of CD3+ and CD19-is a marker of T-cells, while CD3- and CD19+ characterizes B-cells. Here, we use ADT as an independent annotation of our estimated partition, which is solely based on scRNA-seq expression.

Before applying AWST, we retain the cells having at least 500 detected genes. This pre-processing is a typical step in single-cell analysis, in which low-quality samples may be disrupted or dying cells, or empty droplets (Lun *et al*., 2019). The total number of cells is reduced to 7,613 from the original size of 7,985 (about 5% removed). AWST is applied to a data matrix of 7,613 cells measured on 17,014 features after full quantile normalization. The gene-filtering step (with default parameters) returns 230 genes. None of the 230 genes retained by the filtering step is included in the set of antibodies in the ADT data.

Applying hierarchical clustering with Ward’s linkage and Euclidean distance, we obtained a partition of six clusters (Figure 5a), each of which is, for the most part, interpretable according to the levels of expression of the independent ADT data. For instance, cluster CBMC1 identifies the T-cells sub-population given that CD3 is highly expressed, while CD19 is low. The opposite expression of these two markers (CD3 low and CD19 high) identifies the B-cells, clustered in CBMC2. The monocytes are the cells in CBMC5 where the markers CD14 and CD11c are both expressed. Each of the six clusters shows compact sub-groups that are not explained by the ADT, suggesting that the scRNA-seq data may lead to more granularity in the clustering compared to the relatively few ADT markers.

**Figure 5.**
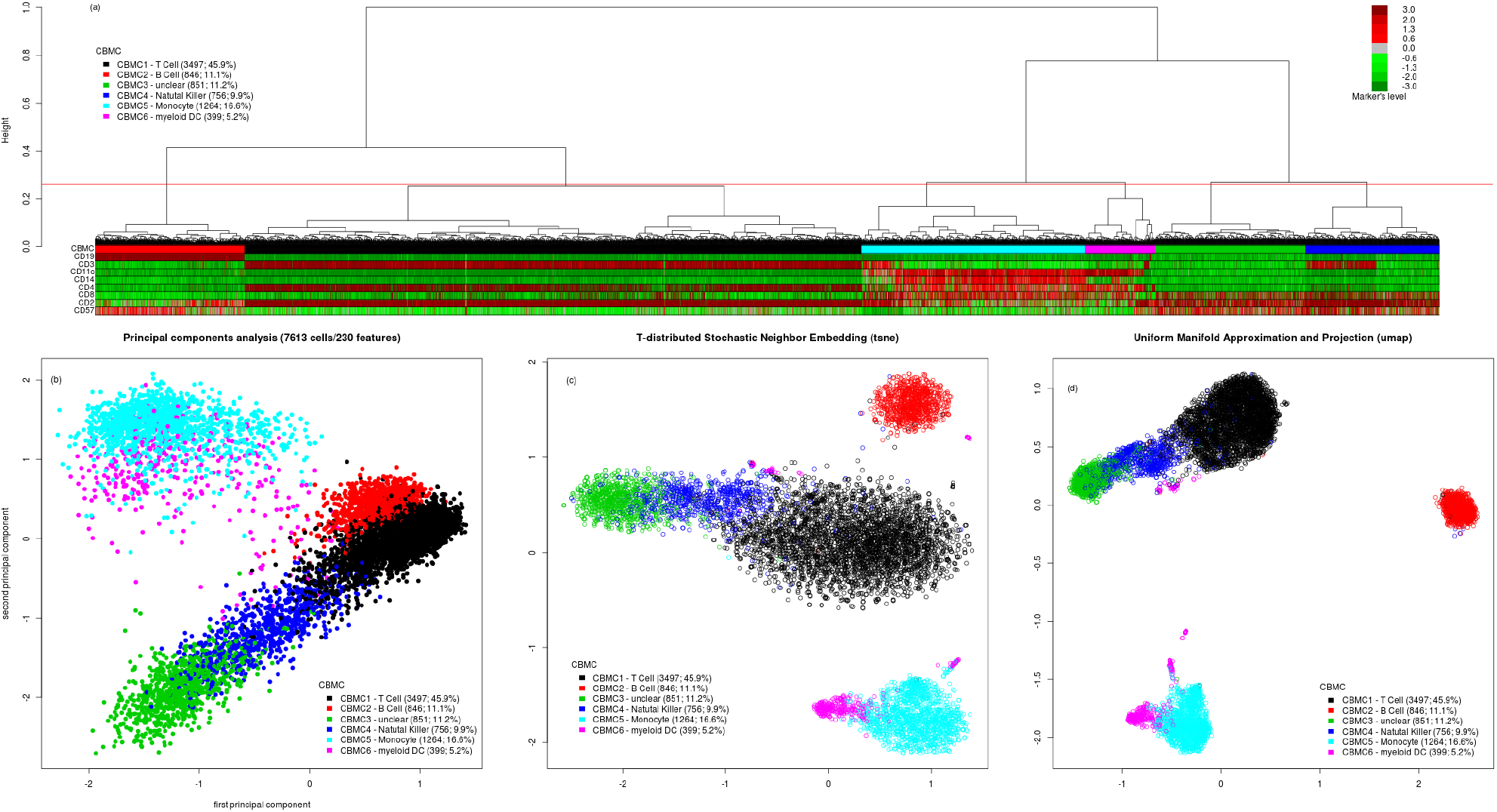
Hierarchical clustering of the CBMC dataset (Stoeckius *et al*., 2017). (a) Hierarchical clustering with Ward’s linkage and Euclidean distance of 7,613 single-cell profiles and 230 features; the first colored bar corresponds to the partition obtained after AWST on the single-cell RNA-seq data; the colored bars below represent the expression of the independent ADT markers; (b) first two principal components annotated with the partition from the hierarchical clustering; (c) t-SNE plot annotated with the partition from the hierarchical clustering; (d) UMAP plot annotated with the partition from the hierarchical clustering.

The inferred clusters are well separated in the space of the first two principal components computed from the 230 features after AWST (Figure 5b). A 3-dimensional view of the cells is at https://tinyurl.com/ybp3wydb. Overall, there is good agreement between ADT data and scRNA-seq-based unsupervised clustering (Supplementary Figures 17 and 18).

Generally, t-distributed Stochastic Neighbor Embedding (t-SNE; figure 5c) mapping (van der Maaten and Hinton, 2008), and Uniform Manifold Approximation and Projection (UMAP; Figure 5d) (McInnes *et al*., 2018) are the preferred visualization methods for single-cell data. The hierarchical clustering of Figure 5a shows its superiority to unveil the complex similarity between cells. From tSNE or UMAP, the landscape of this collection of cells appears quite flat, while the hierarchical clustering is able to unveil the relations between cell types. Overall, this analysis demonstrates that AWST leads to biologically interpretable results with a simple hierarchical clustering procedure in single-cell data.

## 4 Discussion

Data transformations are an important step prior to unsupervised clustering. However, not much attention has been paid to the effects of standardization on downstream results. We have presented a novel data transformation, AWST, with the aim of reducing the noise and regularizing the influence of outlying genes in the clustering of RNA-seq samples.

AWST is a per-sample transformation that comprises two main steps: the standardization step exploits the log-normal distribution to bring the distribution of each sample on a common location and scale; the smoothing step employs an asymmetric winsorization to push the lowly expressed genes to a minimum value and to bound the highly expressed genes to a maximum.

Our optional filtering procedure involves a heterogeneity index, Shannon’s entropy, that does not need a preliminary location estimate, unlike the variance and its robust alternatives. This strategy is possible because of the smoothing step, which maps the values of all samples into the same interval.

Applying our transformation to synthetic and real datasets, including bulk and single-cell RNA-seq experiments, shows that AWST leads to biologically meaningful clusters. We have used synthetic data to compare AWST with two recent protocols, and two more pre-processings, based on the Hart transformation and on FPKM, respectively. AWST yields the best results with any clustering method, while the competitors’ performances depend on the downstream clustering algorithm.

In a LGG real dataset, AWST followed by hierarchical clustering with Ward’s linkage and Euclidean distance uncovers cancer subtypes previously discovered in independent, sometimes multi-omic, experiments.

Applied to single-cell data, AWST leads to compact, biologically meaningful clusters, consistent with independent external data.

Overall, AWST leads to better and more robust sample segregation allowing to move from standard statistical discovery to a more in-depth investigation of sub-groups within clusters on the way to precision medicine. Furthermore, AWST allows to use statistical coherent methods, such as the Wards linkage method applied to Euclidean distance, (both related to the ANOVA setup), which is the only guarantee for the trustworthiness of the results in unsupervised settings.

## Acknowledgments

The authors thank Chiara Romualdi for fruitful discussions and for her feedback on the manuscript.

## Funding

DR was supported by Programma per Giovani Ricercatori Rita Levi Montalcini granted by the Italian Ministry of Education, University, and Research. SMP was supported by the Department of Science and Technology, Università degli Studi del Sannio, Benevento, 82100, Italy, and the AIRC foundation under IG 2018 - ID.21846 project.

## Notes

### Competing Interest Statement

The authors have declared no competing interest.

