## Supplementary Material for "Per-sample standardization and asymmetric winsorization lead to accurate clustering of RNA-seq expression profiles"

June 4, 2020

### 1 Synthetic data experiment

#### 1.1 Data generation

To simulate realistic data, we started from the SEQC dataset and added random perturbations to a subset of genes to obtain differential expression between groups. The SEQC/MAQC-III project (Su *and others*, 2014) performed an extensive RNA-seq experiment on technical replicates of benchmark biological samples. We used only sample A, corresponding to Agilent's Universal Human Reference RNA (UHRR), sequenced at the Mayo Clinic using the Illumina HiSeq200 platform ("ILM_aceview_gene_MAY"). We downloaded the data from the Bioconductor package `seqc` (Liao and Shi, 2019), and obtained the data matrix of the 80 technical replicates of sample A. We only retained the 19,701 features (out of 55,950) with an ENTREZ identifier.

We then created $k = 5$ groups of samples, each made of $M = 30$ replicate samples, randomly selected (with replacement) from the original set of 80. For each group, we randomly selected (without replacement) $J = 500$ genes, whose expression was altered according to the following multiplicative model.

$$\tilde{y}_j = y_j \cdot (0.001 + r_j), \quad j = 1, \dots, J,$$

where $y_j$ denotes the observed expression of gene $j$, $\tilde{y}$ denotes the perturbed expression, and $r_j$ is the realization of a Gamma random variable with shape parameter $a = 0.5$ and scale parameter $s = 1$. The choice of the Gamma distribution is consistent with typical models for RNA-seq data, in which a negative binomial (i.e., a Gamma-Poisson mixture) is assumed as the data-generating distribution.

Since we selected the genes to alter by sampling without replacement, and since we generated 5 groups, in our simulation a total of $2,500$ genes (out of $19,701$ genes) are responsible for the differences among groups, making this a realistic yet challenging dataset for cluster analysis.

#### 1.2 AWST pipeline

We apply AWST (default values) to the raw-counts of the synthetic data. We omit the gene-filtering step to be on the same ground as Hart's transformation. From the Euclidean distances obtained from AWST, we computed the silhouette indexes (Rousseeuw, 1987) (Supplementary Figure 2a). Starting from the same distance matrix, we applied hierarchical clustering with Ward's linkage (Figure 2a and Supplementary Figure 1a). Figure 3a, and Supplementary Figure 3a are the result of ConsensusClusterPlus (Wilkerson *and others*, 2010) applied to the AWST-transformed data, where we required Pearson's correlation as distance and average linkage for the inner and outer hierarchical procedures (default parameters). Supplementary Figure 4a is the result of the ConsensusClusterPlus applied to the AWST-transformed data, requiring Euclidean distance and Partitioning Around Medoids (PAM) clustering (Kaufman and Rousseeuw, 1990).

#### 1.3 Hart transformation

The aim of Hart *and others* (2013) is to divide the features *"into active genes carrying out the work of the cell and other genes that are likely the by-products of biological or experimental noise"* (Hart *and others*, 2013). They do not consider unsupervised clustering as downstream analysis of their data transformation. Nevertheless, we included their contribution because it is close to our proposal. Their protocol starts from FPKM-normalized data. Then,

1) *data are* $\log_2$*-transformed, and*

2) *mean and variance of the transformed values are estimated considering the right tail of the distribution starting from the mode.*

Their final step is

3) *the usual standardization of the* $\log_2$*(FPKM) with the estimates of location and scale obtained in point 2.*

The standardized values are the zFPKM scores. According to their study, the active genes are those having a zFPKM > -3.

The results on the performance of Hart's transformation are in Figure 2b, Supplementary Figure 1b, Supplementary Figure 2b, Supplementary Figure 3b, and Supplementary Figure 4b. Having no indication from Hart *and others* (2013) on how restrict the number of features for downstream analyses, we considered only those genes with an average zFPKM $> -3$. In the synthetic dataset, all the features fulfill this constrain.

### 1.4 TCGA pre-processing and protocol

This protocol is the one defined initially to group the samples in the Lower Grade Glioma study (TCGA Research Network, 2015).

The TCGA protocol comprises these steps

1) *"Gene-level data was restricted to genes expressed in at least 70% of samples."*

2) *"Data were Log2 transformed and median centered across samples."*

3) *"The most variable genes were selected as the 1500 genes with the highest median absolute deviation."*

4) *"Consensus clustering was performed using ConsensusClusterPlus (1000 iterations, resampling rate of 80%, and Pearson correlation)"*

In point 4, average linkage method (default) is supposed as inner and outer linkage.

We refer to steps 1-3 as TCGA pre-processing, and to steps 1-4 (full procedure) as TCGA protocol.

The result of the pre-processing pipeline, paired with Euclidean distance and Ward's linkage for hierarchical clustering, is in Figure 2d (replicated in Supplementary Figure 1d). The silhouette plots of the Euclidean distance matrix with the theoretical partition are shown in Supplementary Figure 2d. Figure 3d (replicated in Supplementary Figure 3d) shows the results of the TCGA protocol. Supplementary Figure 4d shows the results of the TCGA pre-processing followed by ConsensusClusterPlus with Euclidean distance and PAM clustering.

### 1.5 Radovich pre-processing and protocol

Radovich *and others* (2018) adopted the following steps to obtain a transcriptomic clustering of the thymic epithelial tumor for further integration in the multi-omic characterization.

1) *"After restricting to genes with at least 75% non-zero RSEM values,"*

2) *"the genes with the 1000 highest median absolute deviation (MAD) values were chosen. RSEM values identically equal to zero were replaced into smallest non-zero value."*

3) *"Then a log2 transformation was applied and"*

4) *"the values were median centered by gene and divided by MAD expression of each gene."*

The expression levels matrix feeds ConsensusClusterPlus without any specification of the parameters. Hence we assume Pearson's correlation and average linkage (default values) for both inner and outer linkage.

We refer to steps 1-4 as Radovich pre-processing, and to steps 1-4 followed by ConsensusClusterPlus (full procedure) as Radovich protocol.

Applied to the synthetic data, the pre-processing provides the results in Figure 2c, (replicated in Supplementary Figure 1c), and Supplementary Figure 2c. Figure 3c (replicated in Supplementary Figure 3c) shows the results of the Radovich protocol. Supplementary Figure 4c shows the results of Radovich pre-processing followed by ConsensusClusterPlus with Euclidean distance and PAM clustering method.

### 1.6 FPKM pre-processing

To complete this overview of clustering procedures and data transformations, we included one of the simplest pre-processing procedures. Raw-counts were normalized to Fragments Per Kilo-base of exon model per Million reads mapped (FPKM) and log2-transformed. We restricted the features to the top 2,500 or 5,000 according to their standard deviations. Finally, we standardized by features the restricted data matrix. These steps define the FPKM pre-processing of raw-counts.

Supplementary Figure 1e (2500 features) and 1f (5000 features) show the results of hierarchical clustering with Ward's linkage and Euclidean distance applied to log2-FPKMs. Supplementary Figures 3e and 3f show the results of ConsensusClusterPlot with average linkage (inner and outer) paired with Pearson's correlation as distance matrix. Supplementary Figures 4e and 4f show the result of ConsensusClusterPlot with Euclidean distance and PAM. Supplementary Figures 1e and 1f show the silhouette plots computed with the Euclidean distance according to the theoretical partition.

Supplementary Figure 5 shows the dependency of the Adjusted Rand Index on the choice of the number of features retained in the restricted data-matrix. For each $J = 1{,}000, 2{,}000, \ldots, 10{,}000$, we applied hierarchical clustering with Ward's linkage and Euclidean distance. For each clustering we computed the ARI considering the theoretical partition.

### 1.7 Supplementary Figure 1

This figure extends Figure 2, and has been generated with the script SyntheticExperiment.Rmd (@GitHub).

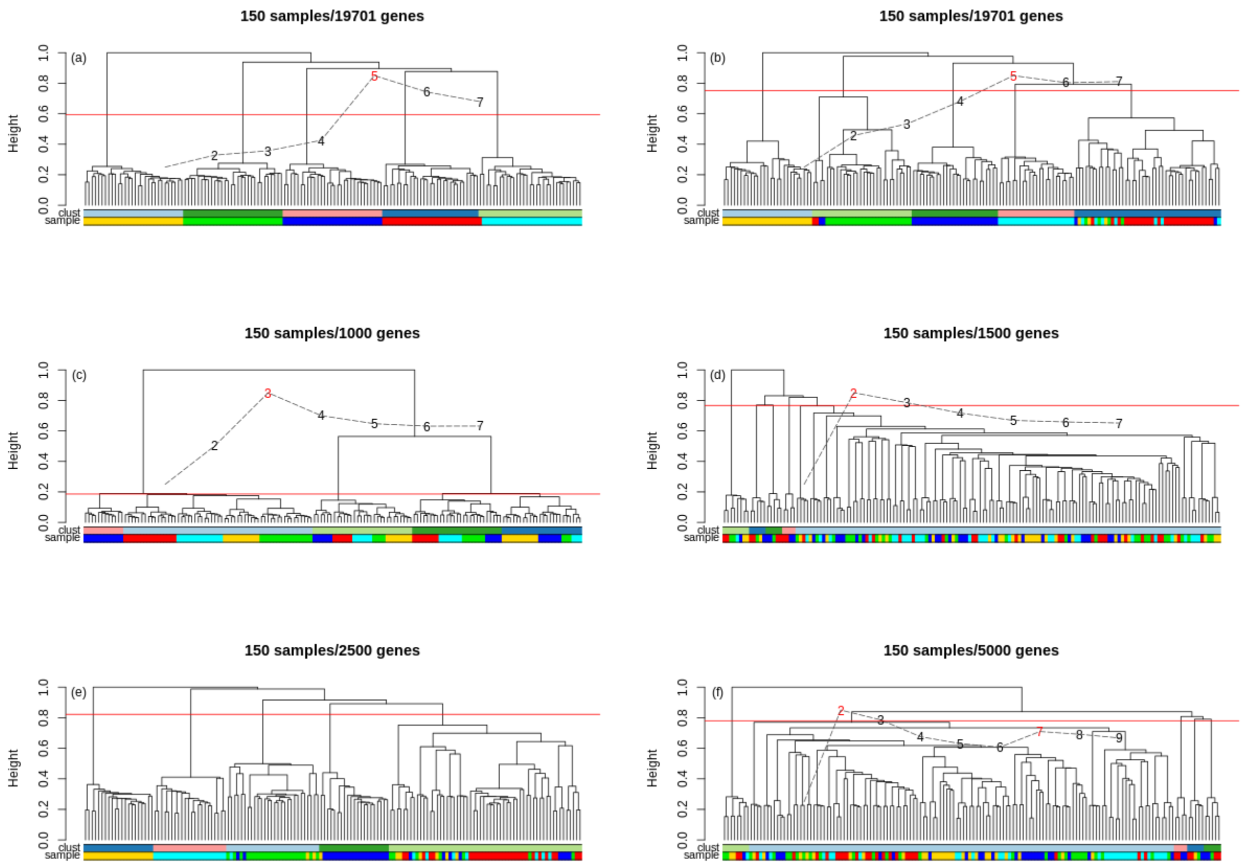


Supplementary Figure 1: (Extended Figure 2) Performance of different data pre-processing paired with hierarchical clustering (Euclidean distance, Ward's linkage). a) AWST data pre-processing; b) Hart's data pre-processing; c) Radovich's data pre-processing; d) TCGA data pre-processing; e) FPKM pre-processing with top 2,500 features according to standard-deviation; f) FPKM pre-processing with top 5,000 features according to standard-deviation. The "sample" bar indicates the true partition, and the "clust" bar indicates the inferred partition obtained by cutting the tree to obtain 5 clusters. The Calinski-Harabasz curve is superimposed in each panel.

### 1.8 Supplementary Figure 2

This figure has been generated with the script SyntheticExperiment.Rmd (@GitHub).

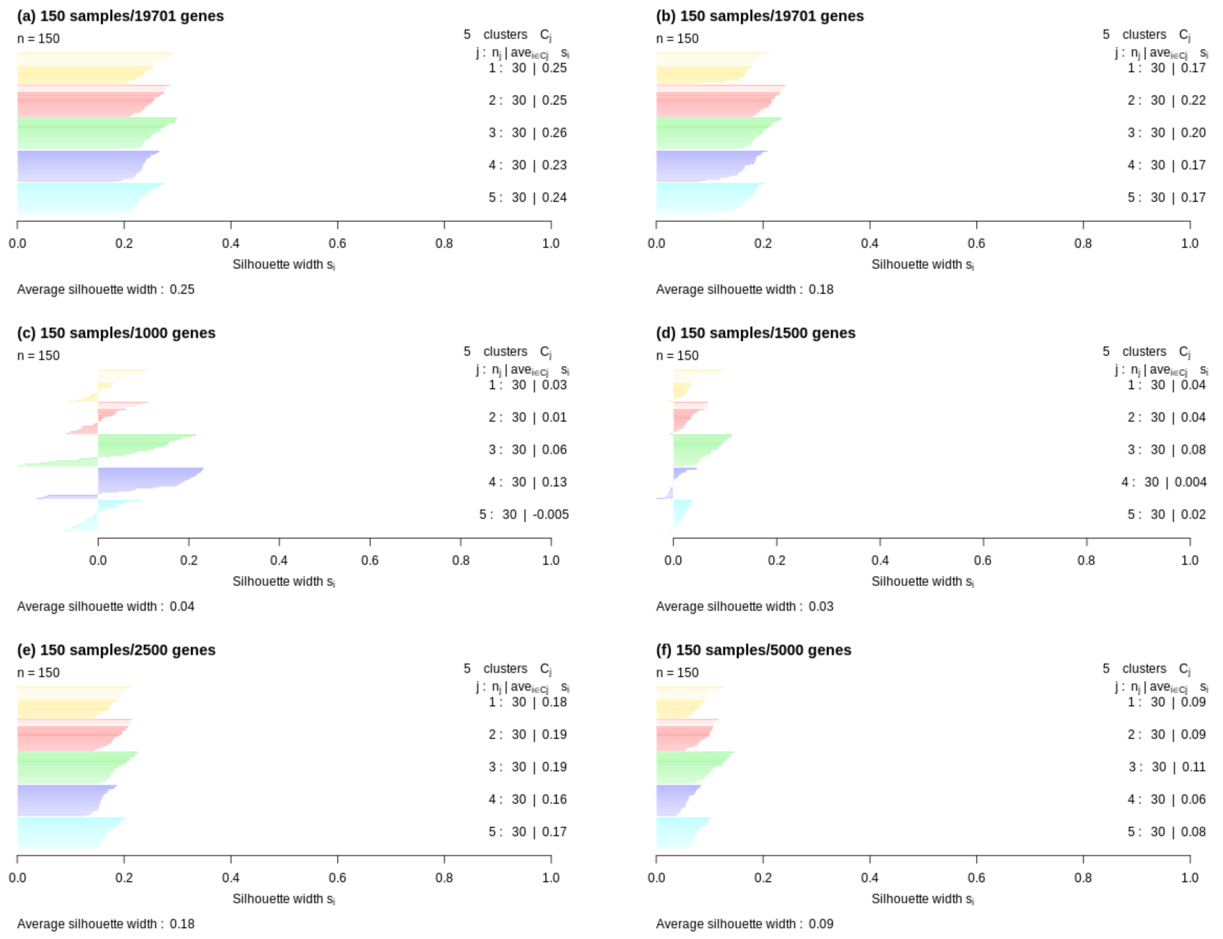


Supplementary Figure 2: Study for the synthetic data of the effects of data pre-processing on the compactness of groups with respect to the theoretical partition and Euclidean distance. a) AWST; b) Hart; c) Radovich;d) TCGA; e) FPKM pre-processing with top 2,500 features according to standard-deviation; f) FPKM pre-processing with top 5,000 features according to standard-deviation.

### 1.9 Supplementary figure 3

This figure extends Figure 3, and has been generated with the script SyntheticExperiment.Rmd (@GitHub).

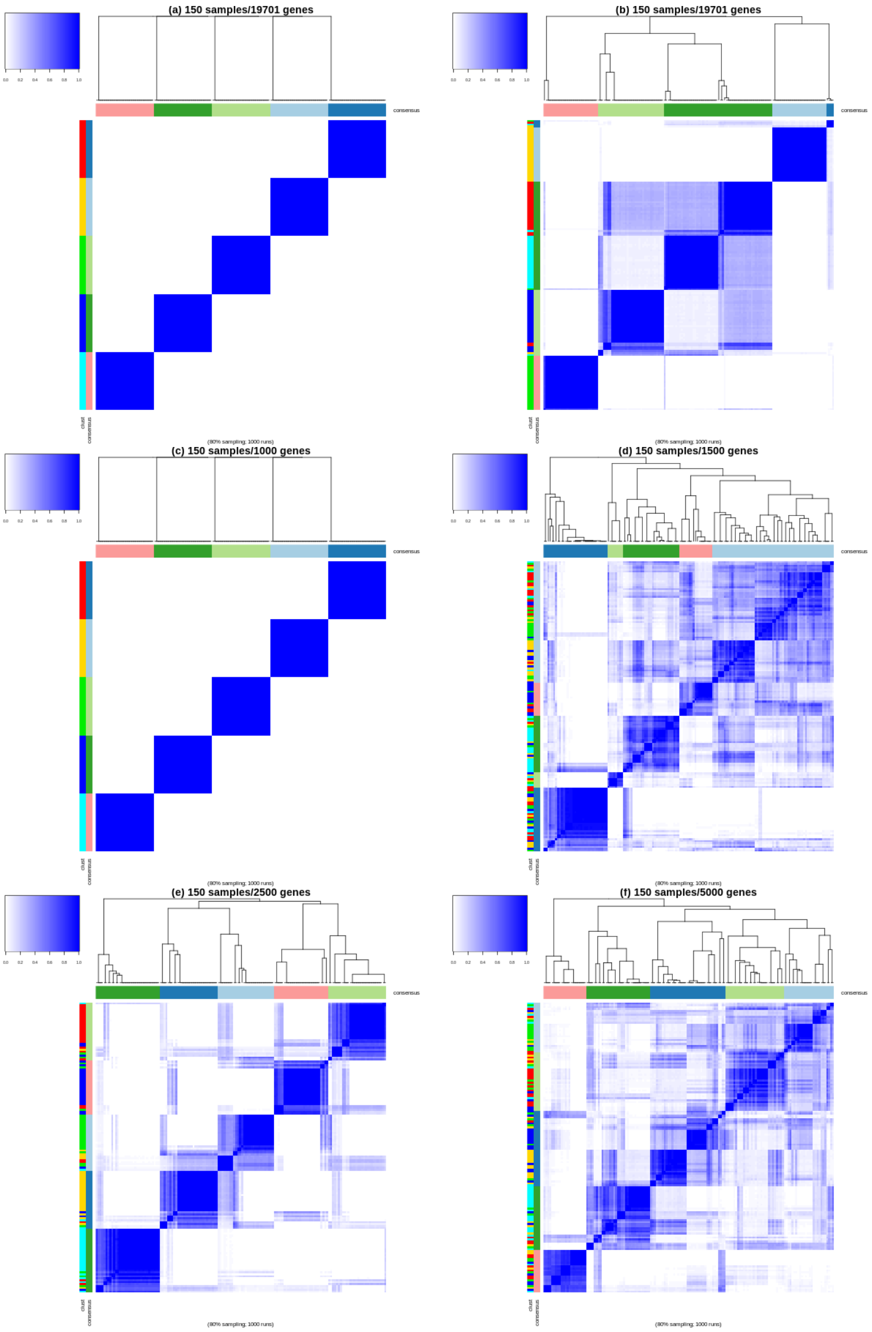


Supplementary Figure 3: (Extended Figure 3) Performance of different data pre-processing paired with ConsensusClusterPlus (inner and outer average linkage with Pearson's correlation as distance matrix). a) AWST data pre-processing; b) Hart's data pre-processing; c) Radovich's protocol; d) TCGA data protocol; e) FPKM pre-processing with top 2,500 features according to standard-deviation; f) FPKM pre-processing with top 5,000 features according to standard-deviation. The "sample" bar indicates the true partition, and the "consensus" bar indicates the inferred partition obtained by requiring 5 clusters.

### 1.10 Supplementary figure 4

This figure has been generated with the script SyntheticExperiment.Rmd (@GitHub).

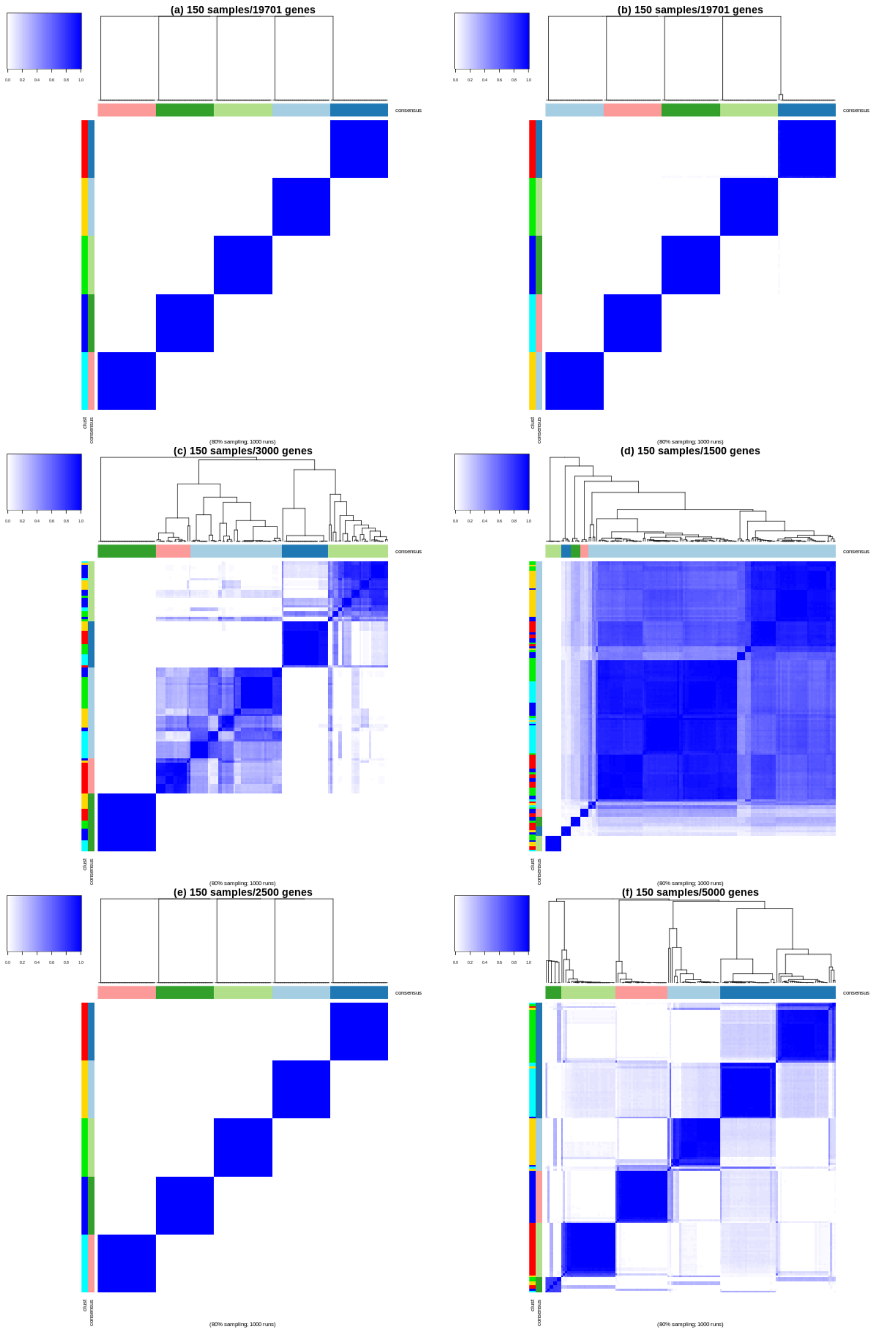


Supplementary Figure 4: Performance of different data pre-processing paired with ConsensusClusterPlus (Euclidean distance and PAM method). a) AWST data pre-processing; b) Hart's data pre-processing; c) Radovich's pre-processing; d) TCGA data pre-processing; e) FPKM pre-processing with top 2,500 features according to standard-deviation; f) FPKM pre-processing with top 5,000 features according to standard-deviation. The "sample" bar indicates the true partition, and the "consensus" bar indicates the inferred partition obtained by requiring 5 clusters.

### 1.11 Supplementary figure 5

This figure has been generated with the script SupplementaryFigure5.Rmd (@GitHub).

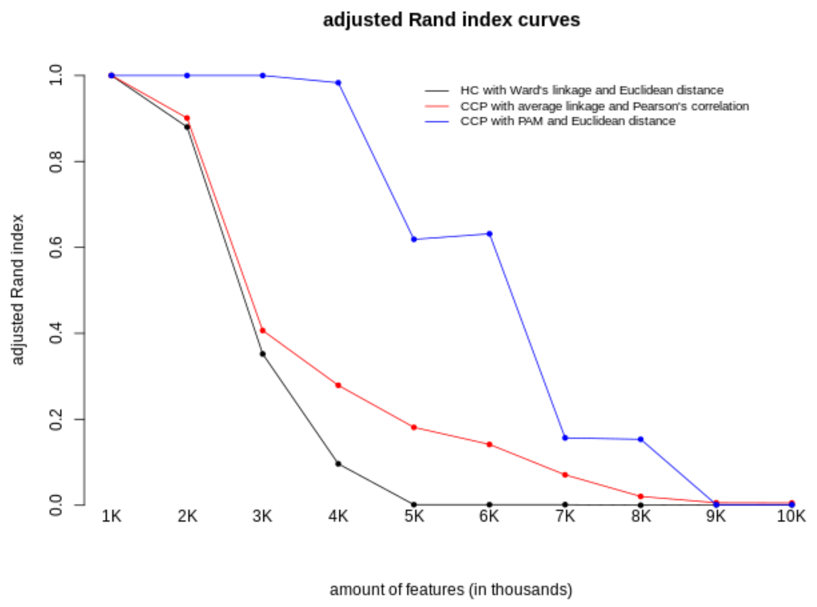


Supplementary Figure 5: Study of FPKM pre-processing on synthetic data. For each amount of features, 1) we applied a clustering procedure, then 2) we computed the Adjusted Rand Index (ARI). Three clustering procedures are shown: black) hierarchical clustering with Ward's linkage and Euclidean distance, red) ConsensusClusterPlus with average linkage (inner and outer) and Pearson's correlation as distance, and blue) ConsensusClusterPlus with PAM and Euclidean distance.

### 2 Lower Grade Glioma study

We retrieved the collection of 516 samples from the TCGA legacy repository. We considered the "level 3" data, specifically the RSEM estimates of 20,531 gene expressions.

We analysed these data by using AWST (Figure 4, Supplementary Figure 11) and Hart (Supplementary Figure 12) transformations, and the TCGA (Supplementary Figures 13 and 14) and Radovich (Supplementary Figures 15 and 16) protocols.

The original study of TCGA Research Network (2015) considered about 230 samples, available at the time.

We applied the TCGA protocol to Lower Grade Glioma study for replicating the original analysis in (TCGA Research Network, 2015) (Supplementary Figure 14 and 15).

Our workflow (with AWST) of the clustering is

1) *correct raw-counts (20,531 features) for GC-content and normalize them with full-quantile as suggested by Risso* and others *(2011).*

2) *Transform the normalized counts (19,138 features) with AWST, and then*

3) *select the (10,212) features with the gene-filtering procedure ($c = 0.01$).* Finally,

4) *apply hierarchical clustering with Euclidean distance and Ward's linkage.*

Steps 1-4 provides the results in Figure 4. Supplementary Figure 11, instead, considers those genes filtered out as non-informative by the filtering step, i.e. those features having a Shannon's entropy less than 0.1.

Clust5 is the partition obtained following the suggestion of the CH-curve. Clust5bis is a second partition in which we explore the compact sub-groups of the clusters in clust5.

In Supplementary Figure 12, we first pre-processed the raw-counts with the Hart transformation, and then we applied hierarchical clustering with the Ward's linkage and Euclidean distance.

### 2.1 Supplementary Figures 6-10

These figures have been generated with the script lgg_awst.Rmd (@GitHub). We used the survival package from Terry M. Therneau and Patricia M. Grambsch (2000).

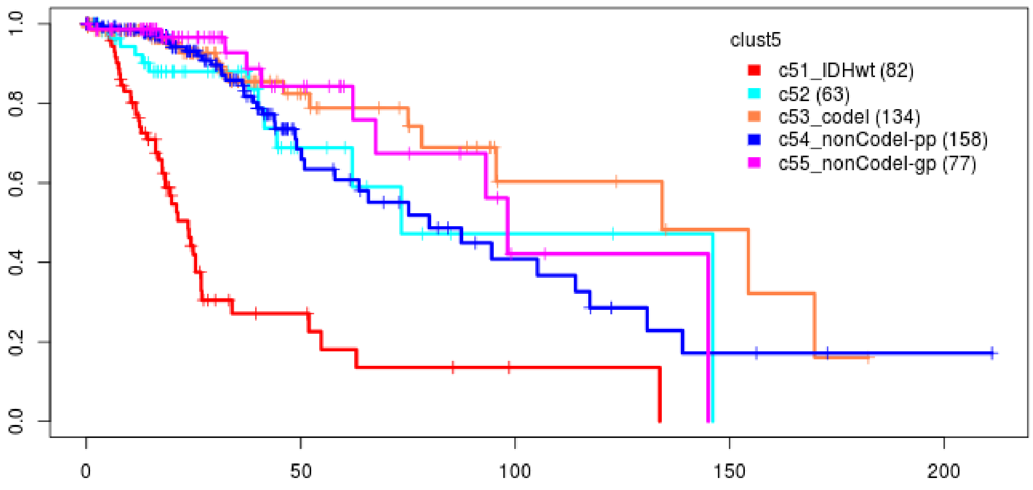


Supplementary Figure 6: Kaplan-Meier estimates of the survival curves associated with clust5 partition, based on the overall survival time. The log-rank test returns *p-value* $\leq 2.2 \cdot 10^{-16}$

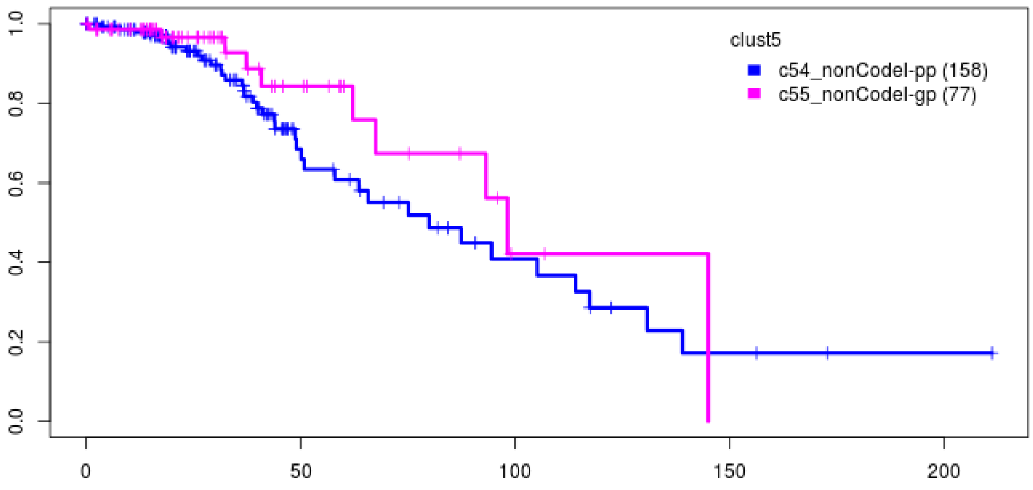


Supplementary Figure 7: Kaplan-Meier estimates of the survival curves associated with the two non-codel subsets c54 and c55, based on the overall survival time. The log-rank test returns *p-value* $= 0.2$.

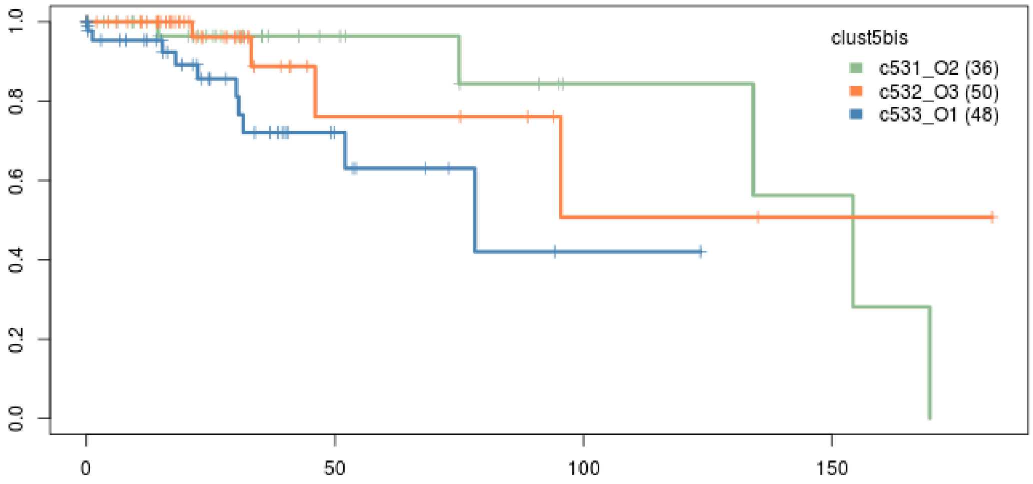


Supplementary Figure 8: Kaplan-Meier estimates of the survival curves associated with the subsets c533/O1, c531/O2, and c532/O3 (Kamoun and others, 2016) in c53 (IDHmut-CODELS), based on the overall survival time. The log-rank test returns $p\text{-}value = 0.02$.

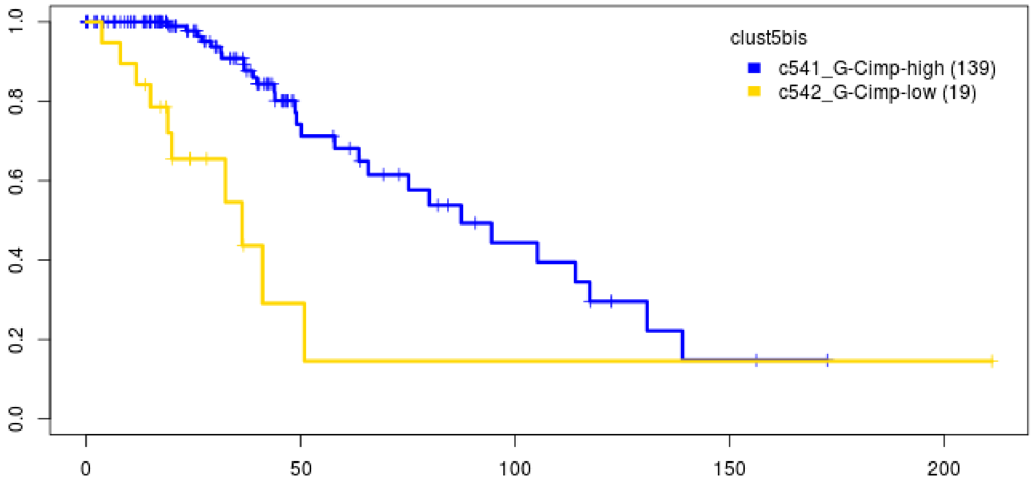


Supplementary Figure 9: Kaplan-Meier estimates of the survival curves associated with the two subsets c541 (G-Cimp-High) ant c542 (G-Cimp-Low), based on the overall survival time. The log-rank test returns $p\text{-}value = 2 \cdot 10^{-4}$.

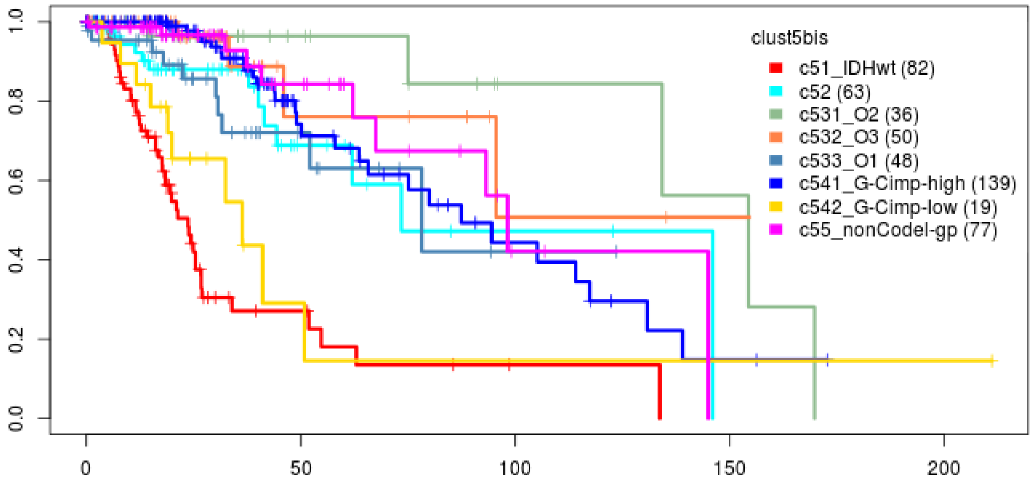


Supplementary Figure 10: Kaplan-Meier estimates of the survival curves associated with clust5bis partition, based on the overall survival time. The log-rank test returns $p\text{-}value \leq 2.2 \cdot 10^{-16}$.

### 2.2 Supplementary Figure 11

This figure has been generated with the script lgg_awst.Rmd (@GitHub).

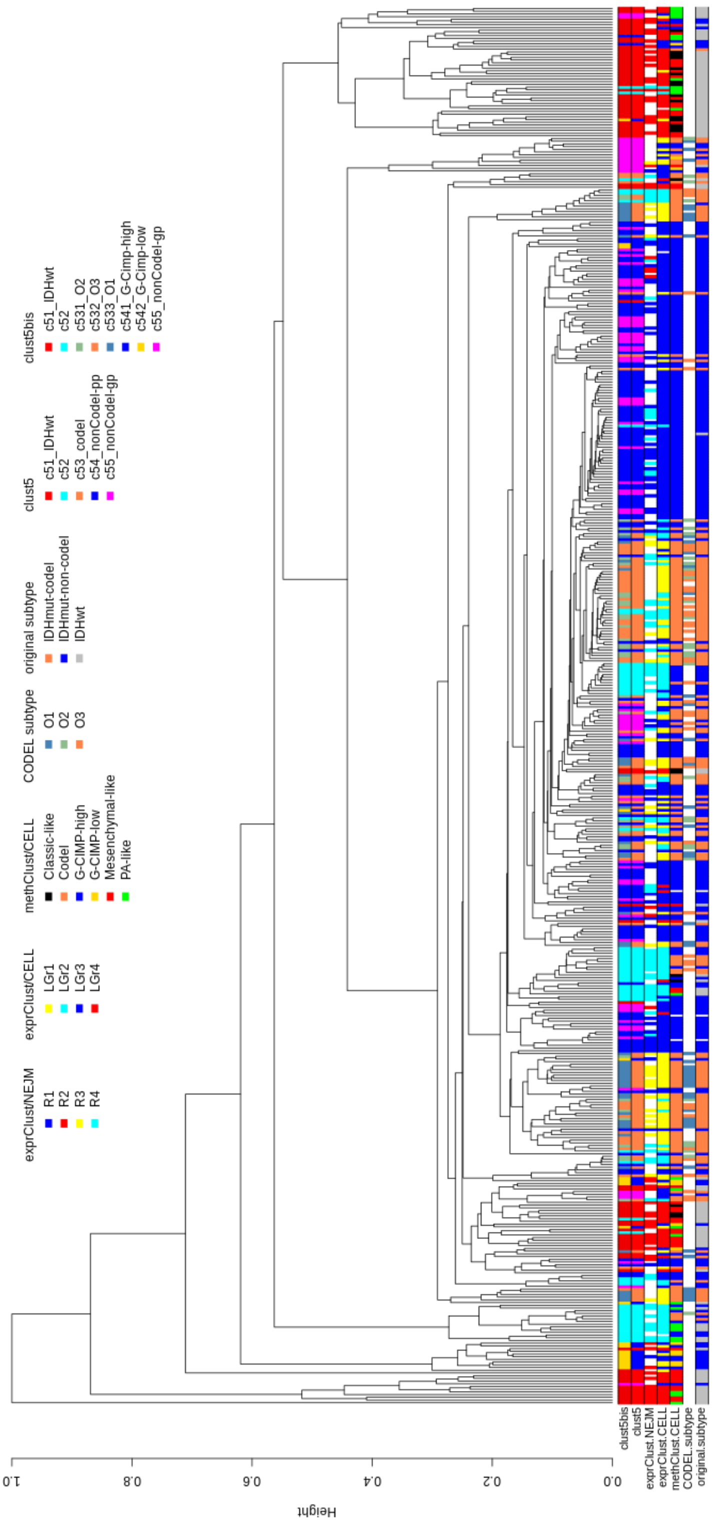


Supplementary Figure 11: Hierarchical clustering of the 516 LGG samples with Wards linkage from and Euclidean matrix involving 8,926 features having heterogeneity less than 0.10 (the features left out by the filtering step).

### 2.3 Supplementary Figure 12

This figure has been generated with the script lgg_Hart.Rmd (@GitHub).

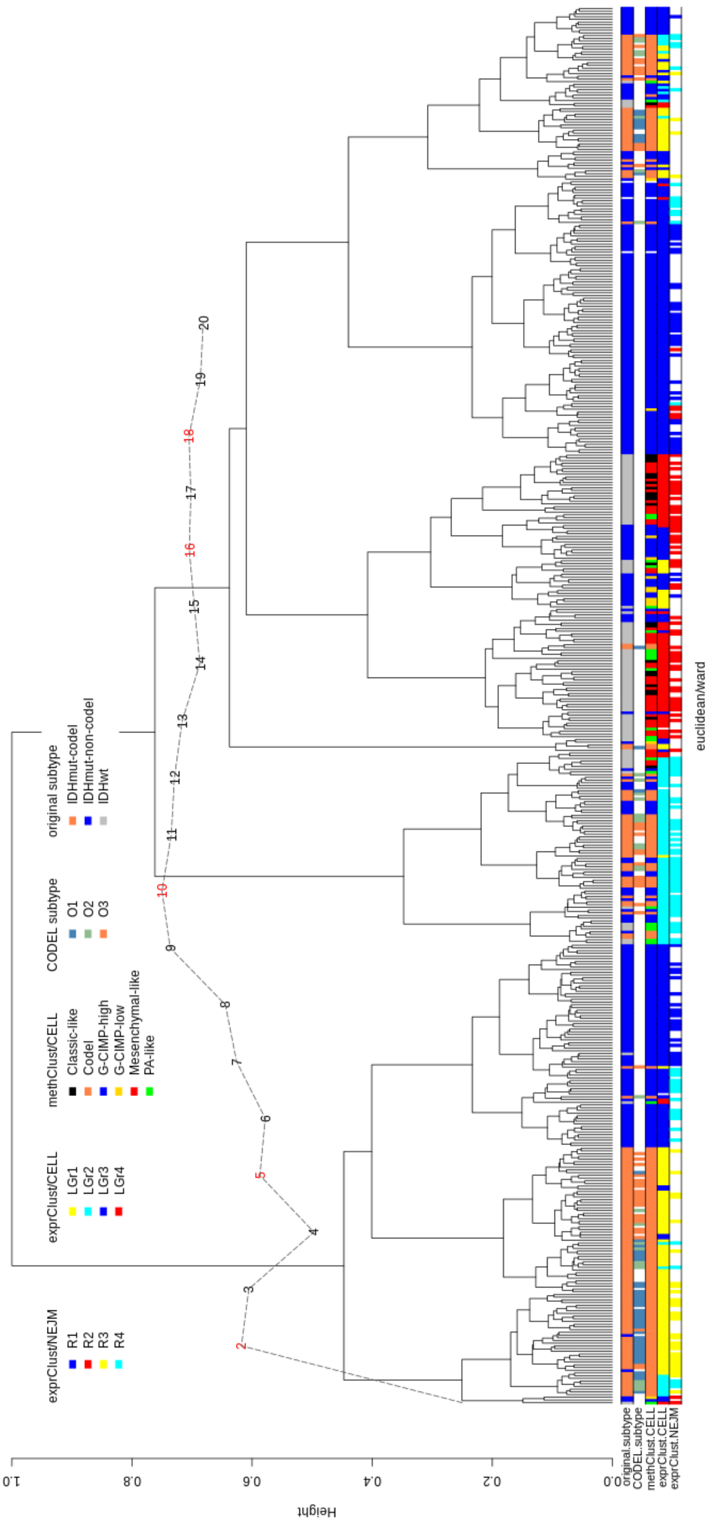


Supplementary Figure 12: Hierarchical clustering of the 516 LGG samples with the Wards linkage method from an Euclidean matrix involving 17,138 features having zFPKM $> -3$ on average (Hart's pre-processing).

### 2.4 Supplementary Figures 13 and 14

These figures have been generated with the script lgg_Radovich.Rmd (@GitHub).

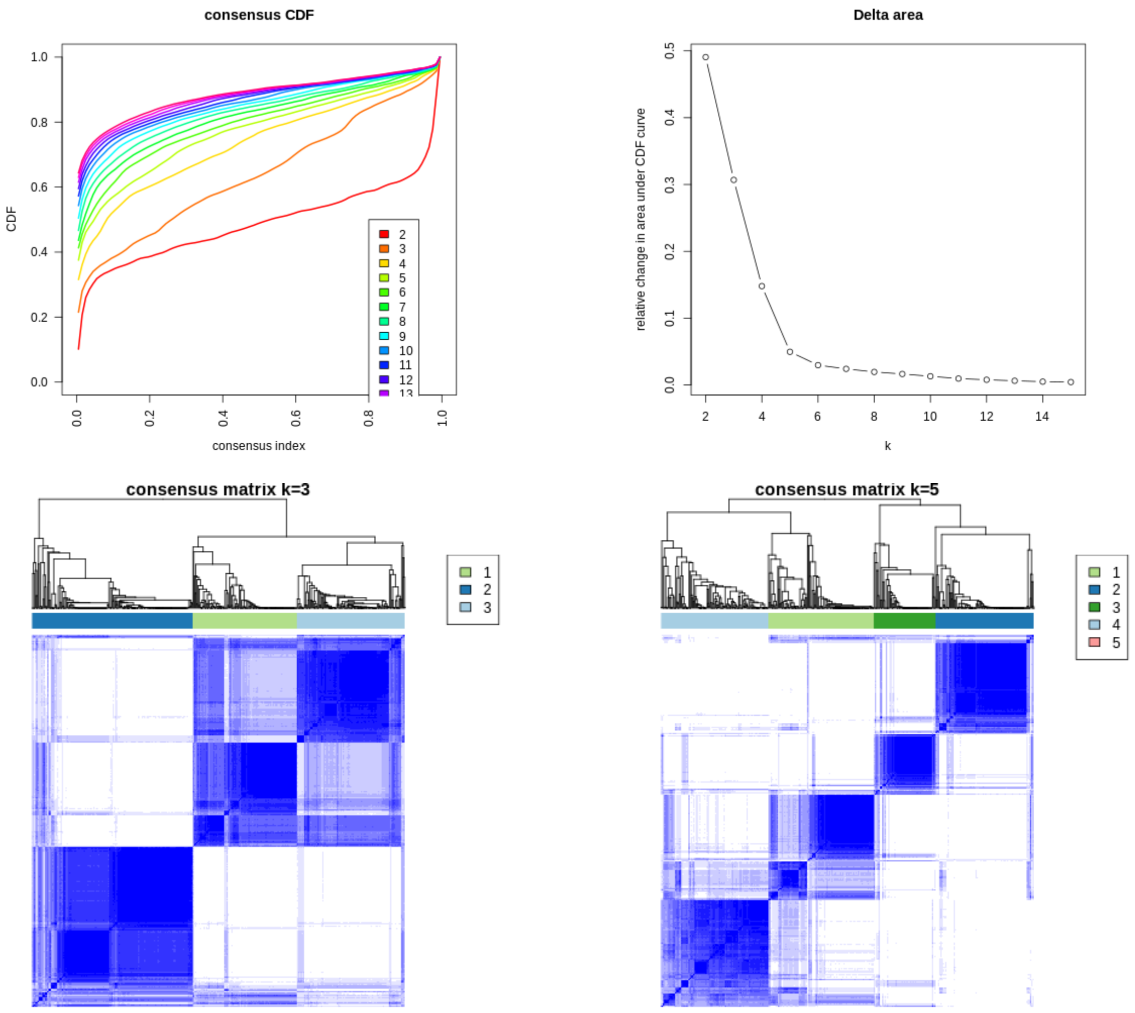


Supplementary Figure 13: ConsensuClusterPlus output (average linkage for inner and outer clustering, and Pearson's correlation matrix as distance) of Radovich's protocol applied to 516 samples of Lower Grade Glioma study.

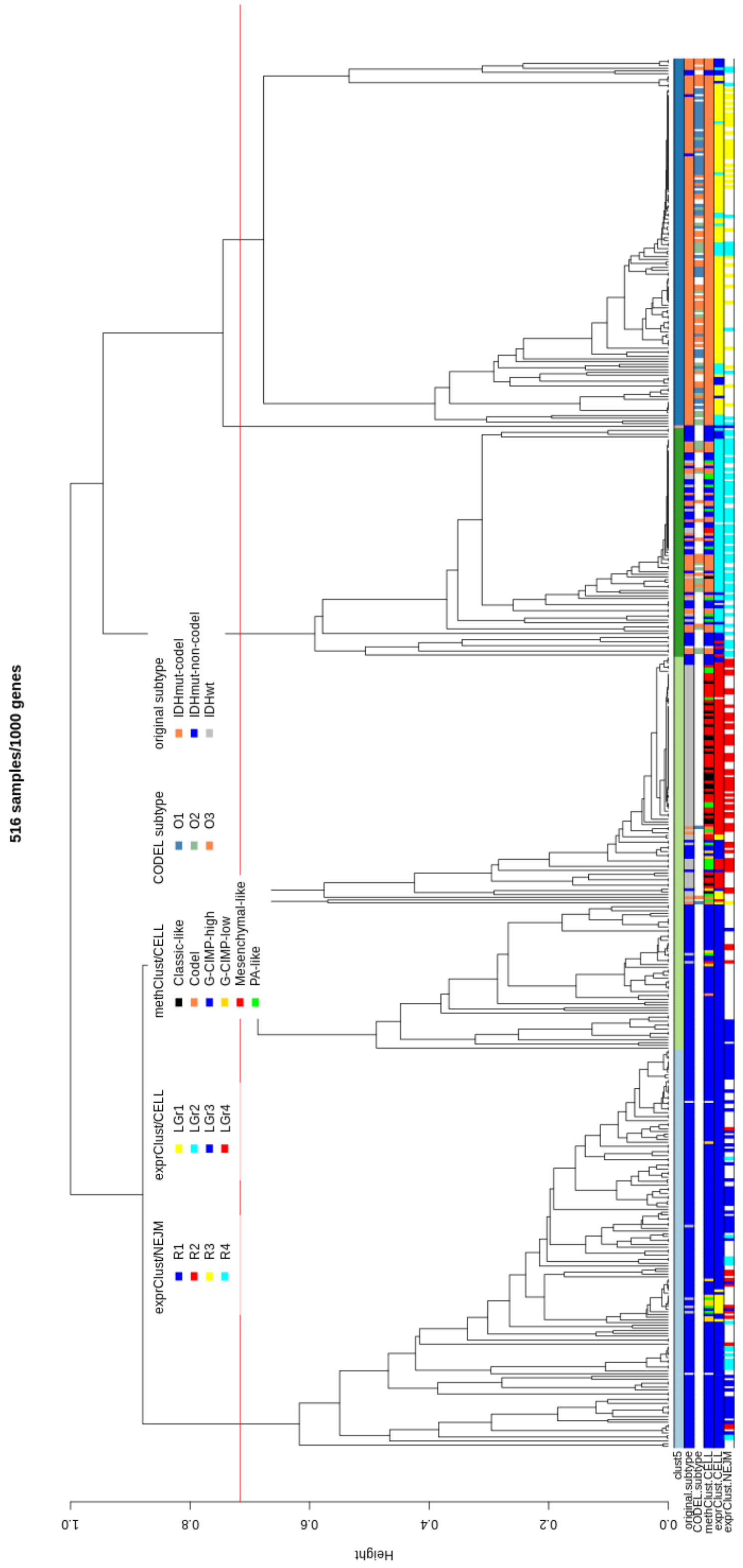


Supplementary Figure 14: Hierarchical clustering of 516 Lower Grade Glioma samples from ConsensuClusterPlus output (average linkage for inner and outer clustering, and Pearson's correlation matrix as distance) in case of five groups for Radovich's protocol.

### 2.5 Supplementary Figures 15 and 16

These figures have been generated with the script lgg_TCGA.Rmd (@GitHub).

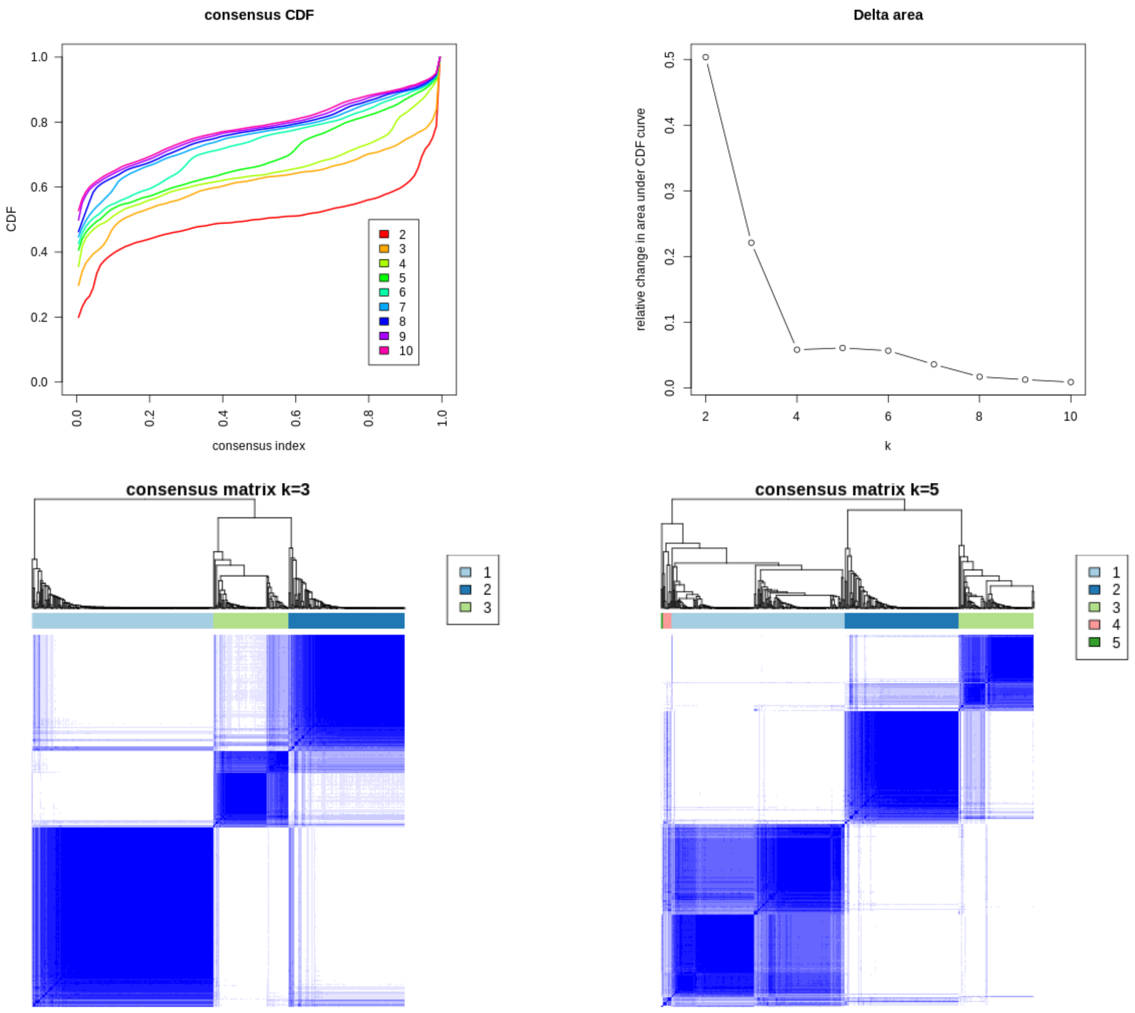


Supplementary Figure 15: ConsensuClusterPlus output (average linkage for inner and outer clustering, and Pearson's correlation matrix as distance) of the TCGA protocol applied to 516 samples of Lower Grade Glioma study

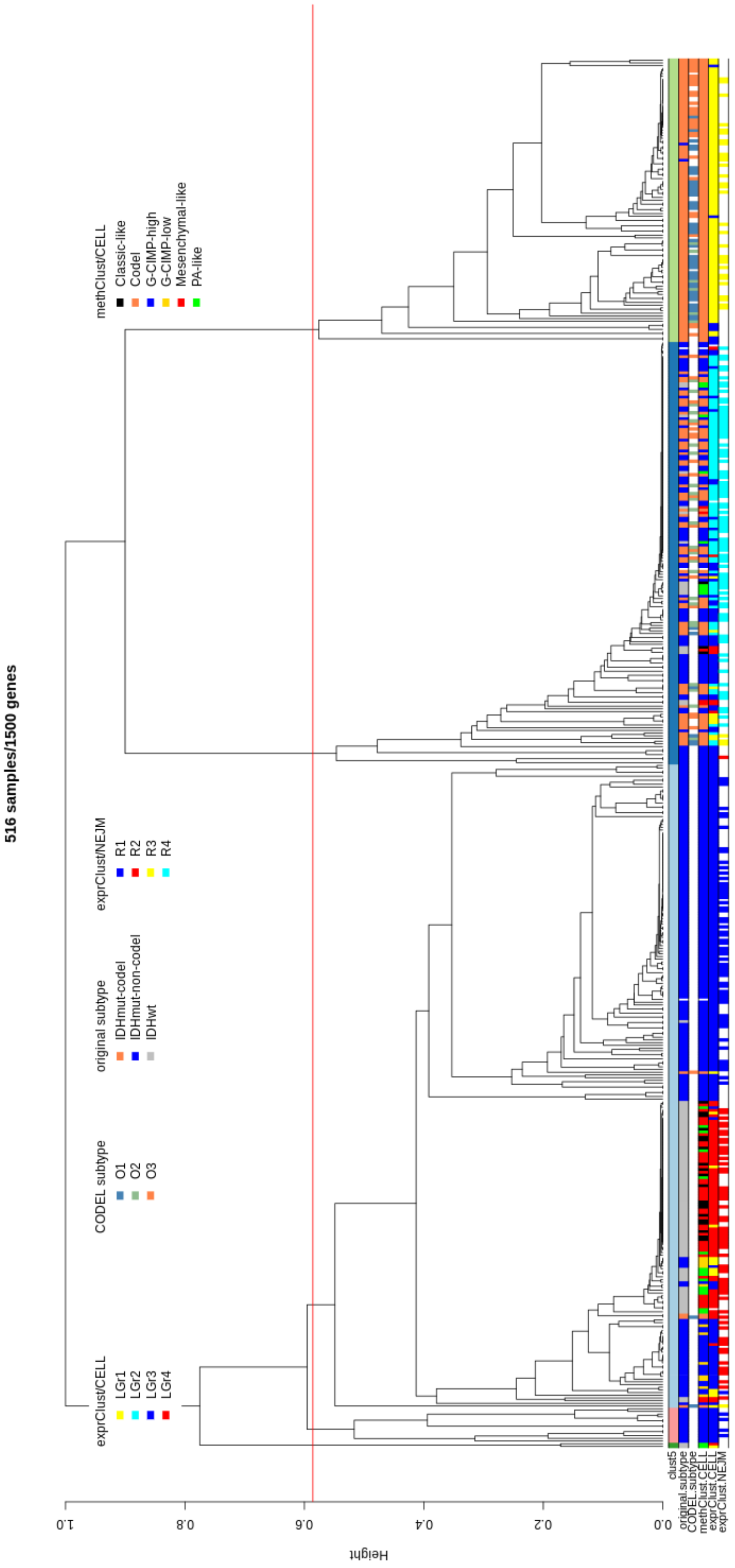


Supplementary Figure 16: Hierarchical clustering of 516 Lower Grade Glioma samples from ConsensuClusterPlus output (average linkage for inner and outer clustering, and Pearson's correlation matrix as distance) in case of five groups for TCGA protocol.

### 3 Cord Blood Mononuclear Cells

We downloaded the data from the Gene Expression Omnibus (GEO) repository, with accession number GSE100866.

The available data include two tables: one with the ADT expression levels and the other with the single-cell RNA-seq expression levels.

To obtain the dendrogram in Figure 5,

1) *we filtered out the samples having a minimum of 500 expressed genes (about the 5% of samples have been removed), then*

2) *we applied the quantile normalization, as implemented the EDASeq R package (Risso* and others*, 2014), and finally*

3) *we applied the AWST transformation with default parameters*

4) *followed by hierarchical clustering with Ward's linkage and Euclidean distance.*

After steps 1-3, we obtained a data-matrix with 7,613 samples (out of 7,985) and 230 features (out of 17,014). From this matrix we computed the principal components (Figure 5b, and the scatter-plots in Supplementary Figures 17 and 18), the two-dimensional t-SNE (Donaldson, 2016) in Figure 5c, and the two-dimensional UMAP (Konopka, 2018) in Figure 5d.

The first three principal components have been arranged to be shown in the TensorFlow embedding projector (Abadi *and others*, 2015) at this link https://tinyurl.com/ybp3wydb. We were not able to control the coloring of cells in the embedding projector, but we fed the application with the cluster codes of CBMC6 partition, as well as the markers for each cell.

### 3.1 Supplementary Figure 17 and 18

These figures have been generated with the script cbmc.Rmd (@GitHub).

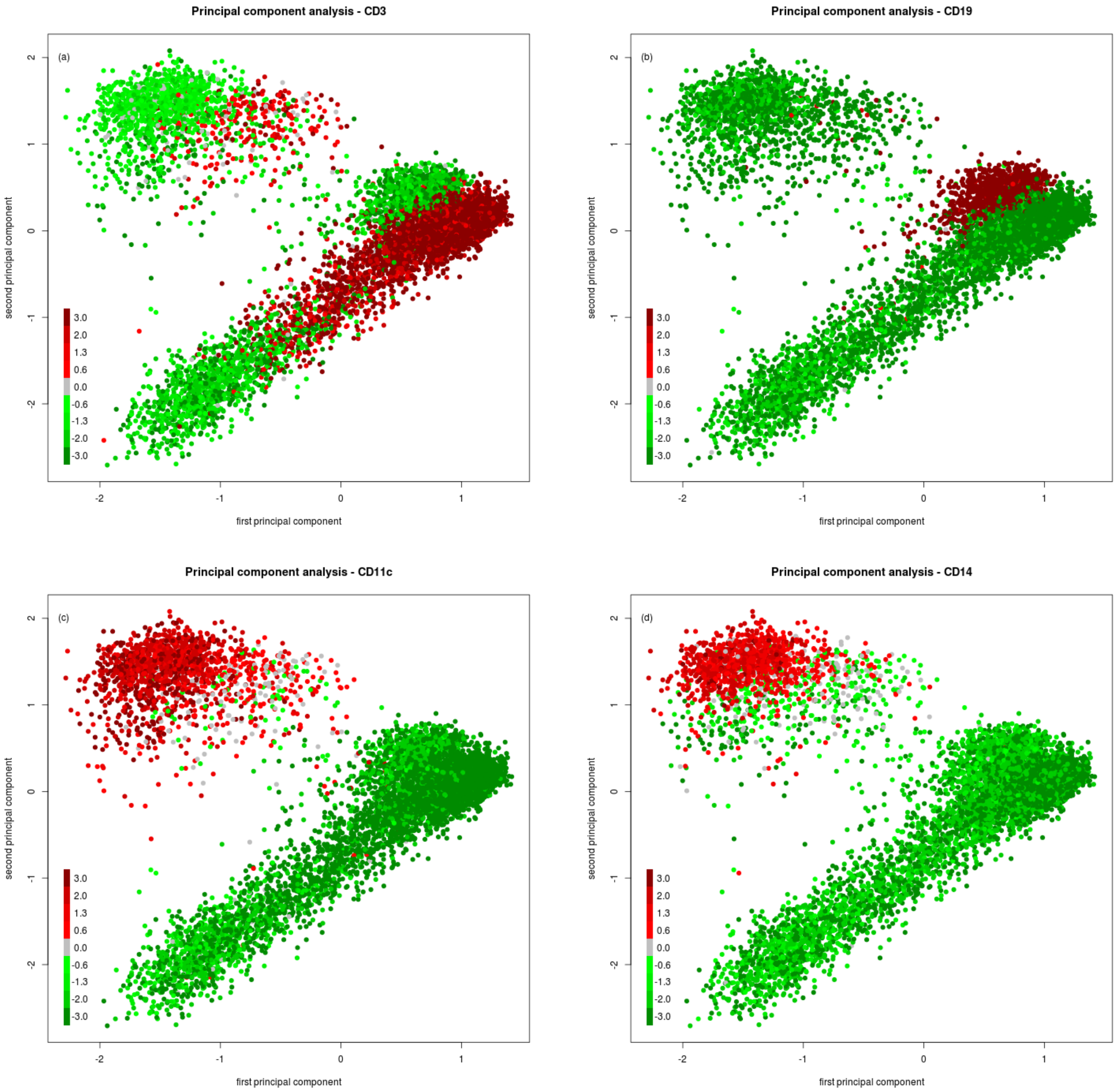


Supplementary Figure 17: Principal components of CBMC computed from AWST data. The color of the cells is the level of expression from the antibody markers independently sequenced. (a) CD3, (b) CD19, (c) CD11c, (d) CD14, (e) CD4, (f) CD8, (g) CD2, and (h) CD57.

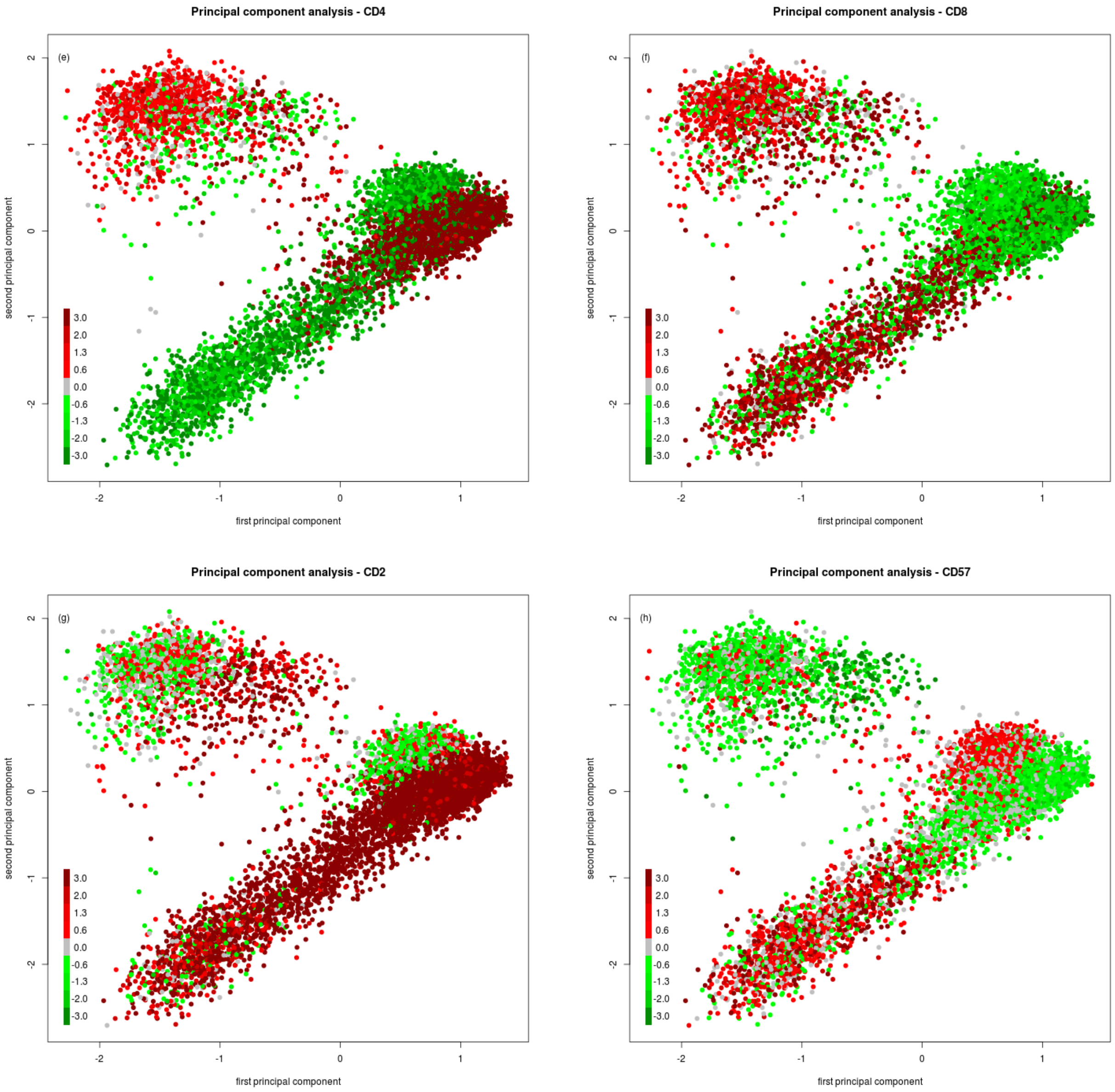


Supplementary Figure 18: Principal components of CBMC computed from AWST data. The color of the cells is the level of expression from the antibody markers independently sequenced. (a) CD3, (b) CD19, (c) CD11c, (d) CD14, (e) CD4, (f) CD8, (g) CD2, and (h) CD57.
